# Past thermal history and heat stress shape patterns of coral-algal symbioses in massive *Porites*

**DOI:** 10.64898/2025.12.02.690324

**Authors:** Lindsey Kraemer, David Juszkiewicz, Zoe T. Richards, Kate M. Quigley

## Abstract

Massive corals, like some *Porites* species, are key reef-builders and provide critical habitat on coral reefs. *Porites* often exhibit high tolerance to stressors, like warming from climate change, although the drivers of these patterns are unclear. Using high-throughput sequencing, we examined whether associations with Symbiodiniaceae underpin this tolerance. Symbiodiniaceae diversity and community composition within two massive species (*Porites lutea* and *P*. *lobata*), whose identities were verified through morphometric examinations, were examined across 200 km of Ningaloo World Heritage Reef, Australia. Contrary to previous studies showing low diversity of high-fidelity symbionts, we found significant variability within *Cladocopium*, which differed significantly across host species, reef sites, and temperature metrics. Locations with broader temperature ranges and a greater number of extreme accumulated heat stress events (measured as ≥ 8 Degree Heating Weeks) exhibited significantly higher prevalence and abundance of thermotolerant *Cladocopium* C116. Community structure also varied significantly by the number of DHW events per location, where a low level of disturbance (1-3 events) resulted in a relatively lower community diversity compared to sites with 3-6 and 6-9 events. The southernmost and coolest site (in terms of Maximum Monthly Mean) harboured the greatest Symbiodiniaceae community diversity, whereas the northernmost and warmest site harboured the greatest Symbiodiniaceae sequence diversity. These lines of evidence suggest that *Porites* may hold greater capacity to vary, and by extension modify, their symbiont community diversity than previously thought. This flexibility could contribute to the genus *Porites* long-term evolutionary success and relative resilience to global marine heatwaves.

## Introduction

Symbiotic relationships are foundational to all ecosystems. Coral reefs exemplify this through the mutualistic interactions between scleractinian reef-building corals and the photosynthetic dinoflagellates of the family Symbiodiniaceae. These algal symbionts live within coral tissue, providing photosynthetically derived energy that supports coral growth and calcification (Muscatine 1990). In exchange, Symbiodiniaceae receive access to essential inorganic nutrients (Muscatine and D’Elia 1978). This symbiosis underpins the ecological and structural foundation of coral reefs, supporting not only the coral hosts and their algal symbionts, but also the trophic levels that depend on reef productivity (Ainsworth et al. 2010).

Climate change is disrupting coral symbiosis through rising sea surface temperature (SST) and frequent accumulated heat stress events, in which SST anomalies exceed the Maximum Monthly Mean (MMM; Strong et al. 1997), often measured as Degree Heating Weeks (DHW; Gleeson and Strong 1995). This leads to coral stress, visualised as bleaching, due to the corals losing their algal symbionts (Sully et al. 2019). Prolonged, extreme temperature conditions can lead to host mortality (Hoegh-Guldberg 1999; Baker et al. 2008). The increasing frequency and intensity of these events, coupled with less stable environmental conditions (Putnam et al. 2017; Sully et al. 2019), pose a significant threat to the persistence of coral reef ecosystems by contributing to the decrease in coral cover globally (Heron et al. 2016; Hughes et al. 2018) and the loss of biodiversity and ecosystem functioning (Wernberg et al. 2024).

Corals can contend with environmental stress, including marine heatwaves, using short-term acclimatory responses or longer-term adaptive responses (reviewed by Voolstra et al. 2021). Host tolerance is influenced by its genetic makeup and the physical tolerance of its prokaryotic and eukaryotic communities (Putnam et al. 2017). Symbiodiniaceae-host associations can be specialised or generalist, where symbionts associate with multiple coral species across various environmental conditions (LaJeunesse et al. 2018). One critical acclimatory mechanism is the change in the relative abundance of Symbiodiniaceae members (shuffling) or the uptake of new symbionts from the environment (switching) (Baker et al. 2004; Quigley et al. 2022). This flexibility in diverse symbiotic associations is thought to aid in withstanding abrupt environmental stress, especially to heat (Buddemeier et al. 2004; Berkelmans and van Oppen 2006; Silverstein et al. 2015; Quigley et al. 2022).

Massive growth forms of the genus *Porites* Link, 1807, can exhibit resistance to environmental stress (Levas et al. 2013). Their stress tolerance is likely partly due to their thick tissue layers, slow skeletal growth, and high investment in energy reserves and cellular repair mechanisms, which together may enhance their ability to withstand thermal fluctuations, physical damage, and disease (reviewed by Putnam et al. 2017), but it is less well known if their Symbiodiniaceae communities contribute to this tolerance. The long-term paradigm is that *Porites* species generally host low-diversity communities of high-fidelity symbionts, particularly within the *Cladocopium* C15 lineage (LaJeunesse et al. 2004; Putnam et al. 2012). However, studies have since revealed that a broader range of symbiont associations may be possible (Ziegler et al. 2015; Pootakham et al. 2018), suggesting that *Porites* species may be more flexible in their associations than initially thought. This flexibility in their Symbiodiniaceae community composition may contribute to their relative resilience compared to other genera, like branching *Acropora* (Depczynski et al. 2013; Gardner et al. 2019), and may be a key factor in their long-term evolutionary success and stress tolerance.

The functional diversity within the Symbiodiniaceae family includes a broad range of physiological responses to environmental stress (Swain et al. 2016; Davies et al. 2023). *Cladocopium* C3, for example, is considered relatively sensitive to stress (Silverstein et al. 2015), whereas *Durusdinium* D1 is generally more stress tolerant, particularly of warming (Swain et al. 2016). However, understanding the functional diversity of Symbiodiniaceae taxa is nuanced, as physiological responses are not strictly genus-specific (Davies et al. 2023) and can vary at the intra-lineage or strain level (Howells et al. 2012; LaJeunesse et al. 2018). Both dominant and background abundances of Symbiodiniaceae are critical to determining host responses to environmental conditions (Quigley et al. 2014). Recent evidence suggests that massive *Porites* can shuffle their Symbiodiniaceae communities in response to heat stress, with some taxa becoming more prevalent following an episode of bleaching-level stress (Starko et al. 2023). Given this new insight, we sought to investigate whether Symbiodiniaceae community composition contributes to the remarkable stress tolerance exhibited by *Porites* along the mid-west coast of Western Australia in the eastern Indian Ocean (Depczynski et al. 2013; Richards et al. 2024). To do this, we examined Symbiodiniaceae community diversity within two *Porites* species within the Ningaloo World Heritage reef using high-throughput sequencing. These insights will be critical to predicting the long-term resilience of these important habitat builders along East Indian Ocean reefs.

## Methods

### 1 Study Design

The study area extended along the Western Australia (WA) coastline from Coral Bay (23°09’15”S 113°45’56”E), to the Muiron Islands (21°40’08”S 114°21’29”E), and within Exmouth Gulf (21°52’53”S 114°11’17”E) (> 200km). Coral collections were made in sheltered lagoonal habitats at seven sites. A total of 118 *Porites* colonies were sampled via snorkel using a hammer and chisel, with ∼ 8-20 colonies sampled per site (see Table 1, Fig. S1). Multiple species of *Porites* (massive growth forms) were sampled at each site, including *Porites lutea* Milne Edwards & Haime, 1851 and *Porites lobata* Dana, 1846. Colonies and their respective corallites were photographed using an Olympus Tough TG-6 underwater camera to aid with ex-situ morphological assessment of each species. All sampled colonies were > 0.5 m in diameter, < 6 m in depth, and > 5 m apart from one another to minimise possible sampling of genetic clones. Fragments (< 40 mm in diameter) were chiselled at the upper-mid section of each colony. This was done to maintain consistency in morphological identification assessment and to minimise possible bias from differences in microbial composition due to intra-colony location (Lewis et al. 2022).

**Table 1.**
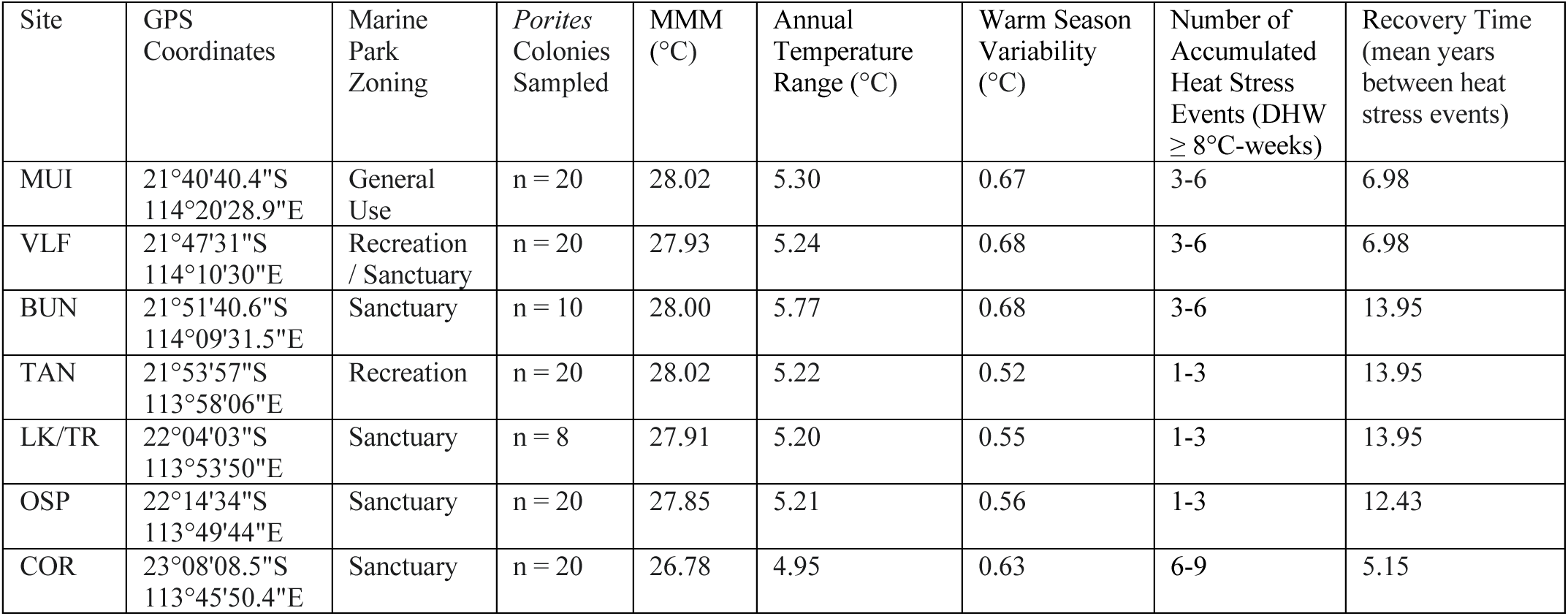
Sampling information and temperature metrics assessed over the climatological period 1985 - 2023. The seven study sites were as follows (listed north to south): Muiron Islands = MUI, VLF Bay = VLF, Bundegi = BUN, Tantabiddi = TAN, Lakeside/Turquoise Bay = LK/TR, Osprey Sanctuary = OSP, Coral Bay = COR.

*Porites* colony fragments were subsampled for 1) taxonomic classification of the coral host and 2) symbiont analysis. Subsamples for host identification (∼ 40 x 40 mm^2^) were placed in bleach for 48 hours, rinsed, and air dried for skeletal examination. Tissue subsamples for symbiont molecular studies were preserved in 100% ethanol and stored at −20°C while at the Minderoo Exmouth Research Lab. To preserve the genetic integrity of samples, ethanol was exchanged within the first 48 hours after collection. Tissue samples were then transported on ice to the Molecular Ecology and Evolution Lab (MEEL) at James Cook University (Townsville, Australia). Upon arrival, samples were topped up with 100% ethanol and stored at −20°C until DNA extractions were performed.

Skeletal fragments of host species were examined using a Leica EZ4W stereo microscope and identified morphologically based on key diagnostic features of the calice, wall, columellae, and septal arrangement (Bernard 1905; Vaughan 1918; Pichon and Veron 1982; Budd et al. 1994). Species-level identification was determined by observations of diagnostic features and morphological descriptions of *Porites* in the Indo-Pacific, particularly those recorded from Ningaloo Reef, WA (Vaughan 1907; Pichon and Veron 1982; Veron 1985, 1986, 1993; Veron and Marsh 1988). Key characteristics used to differentiate *P*. *lutea* and *P*. *lobata* included wall thickness, septal traits (such as the fusion of the ventral triplet, length, and the number of denticles per septum), as well as the height of the pali and the morphology of the columellae (see Fig. S2). After specimen identification, 44 samples were chosen for DNA extraction (Table S1). These samples included fragments identified as *P*. *lutea* (n = 24) and *P*. *lobata* (n = 13), as well as fragments that were uncertain species (potentially *P*. *lobata* or *P*. *lutea*) – requiring further analysis due to the nature of the skeletal sample collected (n = 7).

### 2. DNA Extraction, PCR Amplification, and Sequencing of the ITS2 Region

DNA was extracted from 44 samples using the DNeasy Blood & Tissue Kit (Qiagen, USA) following the manufacturer’s protocol for “Purification of Total DNA from Animal Tissues (Spin-Column Protocol)” with minor modifications (for full protocol see Data S1). PCR amplification was carried out using the multi-copy internal transcribed spacer 2 (ITS2) rDNA region, the most widely used molecular marker to resolve Symbiodiniaceae taxonomy (Davies et al. 2023). The ITS2 region (∼350 base pair (bp)) was amplified from the chosen 44 samples using Symbiodiniaceae-specific primers, ITS-DINO (Pochon et al. 2001) and ITS2Rev2 (Stat et al. 2009). PCR amplification was performed using procedures previously described by Quigley et al. (2022), as detailed below.

Initial PCR amplification was conducted using MyTaq DNA Polymerase (Bioline Inc.) to ensure reliable amplification of the ITS2 region from selected DNA template (see Data S2 for protocol). For high-throughput sequencing, the ITS2 region was amplified using AmpliTaq Gold 360 Master Mix (Thermo Fisher Scientific, USA). Each 25 µl PCR included: 12.5 µl 1x AmpliTaq Gold 360 Master Mix, 1 µl each 0.4µM Forward (5’- GTGAATTGCAGAACTCCGTG - 3’) and Reverse (5’-CCTCCGCTTACTTATATGCTT - 3’) primers, 9.5 µl MilliQ water, and 1 µl DNA template. PCR amplification cycles were performed on an Eppendorf Mastercycler^®^ nexus GSX1 as follows: 95°C for 7 min; followed by 32 cycles at 95°C for 30 s, 57°C for 30 s, 72°C for 30 s, and 72°C for 7 min. PCR products were visualised on 1.5% agarose gels. For PCR products that did not produce a visible band, template dilutions were performed (Table S1), and PCR was repeated until a clear band was confirmed by gel electrophoresis.

Library preparation and next-generation sequencing of ITS2 amplicons were completed by GenomicsWA (Perth, Western Australia). Library preparation incorporated universal Illumina adapters (Forward: 5’ – TCGTCGGCAGCGTCAGATGTGTATAAGAGACAG - 3’; Reverse: 5’ - GTCTCGTGGGCTCGGAGATGTGTATAAGAGACAG - 3’) and PhiX genome spiking (30%). Prepared libraries were sequenced on an Illumina MiSeq platform using MiSeq Reagent Kit v2 (2 x 250 paired-end reads).

### 3. Bioinformatics

Sequenced reads were demultiplexed using provided index information and Illumina’s bcl2fastq program (Illumina 2017). The BBTools program ‘BBduk’ (http://sourceforge.net/projects/bbmap/) was used to remove index sequences and residual Illumina adapter content. FastQC and MultiQC were used to assess sequence quality and confirm the absence of adapter contamination. Following this, all downstream analyses were performed in R (version 4.1.2; R Core Team 2021).

Amplicon analysis is necessary for removal of errors introduced during high-throughput sequencing. The multi-copy nature of the ITS2 region means that single nucleotide differences can be biologically significant or genetically trivial when resolving taxonomic boundaries and assessing ecological functions (Quigley et al. 2014; Davies et al. 2023). Clustering sequences based on similarity thresholds can limit noise introduced by intragenomic variants (Arif et al. 2014; Ziegler et al. 2018) and inform functional diversity metrics by mapping to known ancestral haplotypes and sub-types (Correa and Baker 2009; Hume et al. 2019); however, this also means that taxonomic resolution can be coarse, potentially leading to underestimates of biodiversity (Cunning et al. 2017). Hence, for exploring potentially novel diversity, it is critical to employ high specificity during the denoising process to aid in taxonomic delineation and abundance estimates. The “DADA2” package uses a statistical model-based approach in error detection that allows for improved resolution of sequence variants relative to that of OTU-clustering methods (Callahan et al. 2016).

A standard amplicon analysis workflow was performed using “DADA2” (version 1.22.0; Callahan et al. 2016) with modifications specifically for Symbiodiniaceae (Quigley et al. 2019). Briefly, raw (demultiplexed, adapter-trimmed) reads were first filtered based on read quality scores (Phred score > 30), and then by the “DADA2” error modelling algorithm. Forward and reverse reads were then merged, filtered, and amplicon sequence variants (ASVs) were inferred. Although “DADA2” promotes inclusion of subtle yet potentially meaningful genomic differences, it is still crucial to check for possible overestimation of biodiversity. The “LULU” pipeline (version 0.1.0; (Frøslev et al. 2017) collapses any intragenomic variants that are likely to represent the same ‘genotype,’ if present. “LULU” was run on curated ASVs, using both default (95%, 84%) and conservative (70%, 84%) threshold criteria for sample co-occurrence and sequence similarity, respectively (Quigley et al. 2019).

Biologically relevant variants were then taxonomically classified using the RDP Naïve Bayesian Classifier algorithm (Wang et al. 2007) implemented in “DADA2” aligned to a custom database of ITS2 sequences linked to known taxonomy (Arif et al. 2014). A separate blast of curated ASVs was run against the reference database to ensure that identities matched. For variants that did not map to the reference database, a nucleotide BLAST search was conducted using the BLAST+ 2.15.0 suite against the NCBI nucleotide collection (nt) database (Table S2; Camacho et al. 2009). The search parameters were set to an expected value (E value) threshold of 1e-5, match/mismatch of 1, −2, and gapcosts of 0, 2.5 to ensure high confidence in alignments (hits).

### 4. Read Depth, Intragenomic Variation, and Taxonomic Assignment

A total of 19,075,730 sequenced reads were recovered from the 44 samples sent for Illumina sequencing. The mean sequencing depth was 216,770 ± 3,618 (SE) reads per sample. Pre-processing steps removed 0.1% of reads, leaving 18,901,656 reads for input into “DADA2” for amplicon analysis (mean 211,304 ± 5,0178 SE reads per sample). During chimera identification in the “DADA2” workflow, 2,304 of the 3,489 inferred sequence variants (within the expected range of 294 to 304 bp in length) were identified as chimeras. After bioinformatic processing and chimera removal through the “DADA2” pipeline, a total of 8,065,014 paired reads (94.7%; mean 183,296 ± 4,525 SE reads per sample) mapped to the remaining 1,185 ASVs across all 44 *Porites* samples (Table S1). The “LULU” algorithm did not collapse and merge any ASVs in this dataset in either case where co-occurrence and sequence similarity thresholds were set at default criteria (95% and 84%, respectively) or conservative criteria (70% and 84%, respectively). Hence, previous filtering steps had identified and removed intragenomic variants. Given this, the full, cleaned dataset of 1,185 ASVs was retained and imported into “phyloseq” (version 1.46.0; McMurdie and Holmes 2013) for further analysis.

156 of 1,185 ASVs (13.2%) were not mapped to known ITS2 taxonomic identifications within the reference database. Our BLAST search yielded more than 300 significant hits for each of the 156 unclassified variants. All hits were within the family Symbiodiniaceae. Sub-genus level classification was determined based on the accession match with the highest score for percent identity (≥ 98%) and lowest E value (minimum E value: 1e-140; Table S2). In cases of multiple hits to sub-genus taxa with the same score and E value, classified ASVs were named in the order of the appearance of hits (i.e. “C91_C15**”).** Thus, all 1,185 ASVs retrieved from this dataset were identified using the reference database or manual BLAST searches.

Further analysis included only samples morphologically identified as *P*. *lutea* (n = 24) and *P*. *lobata* (n = 13). There was a total of 6,833,662 total reads mapping to 1,001 ASVs remaining within these 37 samples. On average, sequencing depth was 184,694 ± 4,947 (SE) reads per sample, ranging from a minimum of 110,476 reads to a maximum of 267,804 reads (Table S3). Rarefaction curves of untransformed data were assessed to ensure sequencing depth was sufficient to capture ASV diversity across samples (Fig. S3). Rarefaction curves were generated using the ‘rarecurv’ function from the “vegan” package (version 2.6-4; Oksanen et al. 2022).

### 5. Temperature Metrics

To examine the relationship between Symbiodiniaceae community composition and historical temperature metrics, sea surface temperature (SST) data were sourced from the NOAA Coral Reef Watch (CRW) Climatology and Stress Frequency databases (version 3.5; Heron et al. 2014; Skirving et al. 2020). For each site, data were extracted for the following temperature metrics over the climatological period 1985-2023. Maximum Monthly Mean (MMM), defined as the long-term mean SST for the warmest month of the year (°C), reflects the upper boundary of typical thermal conditions that corals historically experience at a given location. Annual temperature range (°C; hereafter referred to as temperature range), defined as the difference between the warmest and coolest monthly mean SSTs, provides an estimate of the typical thermal range experienced at each site. Warm season variability (°C), defined as the interannual standard deviation of daily SSTs over the three-month period encompassing the climatological warmest month, captures year-to-year fluctuations in peak seasonal temperatures. The number of accumulated heat stress events, defined as the summed number of severe bleaching-level heat stress events with DHW ≥ 8°C-weeks, quantifies the frequency with which sites experienced extreme episodes of thermal disturbance over the climatological period (1985-2023). The final metric derived, recovery time, captures the mean time interval (in years) between episodes of severe bleaching-level heat stress. These temperature metrics were derived from 5 km-resolution satellite time series products in NetCDF4 format (NOAA Coral Reef Watch 2024). Data were processed and visualised in R (version 4.1.2; R Core Team 2021) using the “ggplot2” (version 3.5.1; Wickham 2016), “ggspatial” (version 1.1.9, Dunnington et al. 2023), and “sf” (version 1.0.16; Pebesma 2018) packages.

### 6. Statistical Analyses

ASVs, sample metadata, and taxonomic classifications were imported into “phyloseq” to test for patterns within and across Symbiodiniaceae communities from the *Porites* samples chosen for further analysis (n = 37, Table S3). The “ape” package (version 5.8; Paradis et al. 2004) was used to construct a phylogenetic tree so that ASV phylogenetic relationships could be incorporated into downstream analysis. Normalisation and transformation were performed on raw reads using tools from the “DESeq2” package (version 1.42.1; Love et al. 2014). Standard deviation plots were used to evaluate the most appropriate transformation method (Anderson et al. 2006). Based on these assessments, variance stabilising transformation (VST) was selected as the best fit for this dataset (Fig. S4). All analyses, except for alpha diversity, used VST-normalised, abundance data.

To account for both abundance data and phylogenetic relationships between ASVs, weighted UniFrac distances were generated from VST-transformed read counts. These distances represent beta diversity measures, quantifying differences in Symbiodiniaceae community composition between samples or groups, where groups were defined based on tested factors (i.e. host species, site, temperature metrics) (Lozupone et al. 2007). A non-metric multidimensional scaling (NMDS) ordination was performed on the resulting distance matrix to visualise differences in Symbiodiniaceae communities across tested factors (host species, site, and temperature metrics; see Fig. S5 for stress plot). To assess whether dispersion (within-group variation) of symbiont communities differed across factors, we applied a Homogeneity of Dispersion analysis (PERMDISP) using the ‘betadisper’ and ‘permutest’ functions in the “vegan” package (Table S4). The ‘betadisper’ function was used to calculate the distance from each sample to its group centroid, while the permutation-based ANOVA (‘permutest’) tested whether the mean distances to centroids differed significantly between groups. Given uneven sample sizes among host species across sites, we applied the bias.adjust = TRUE parameter to account for potential bias in centroid estimation. Finally, to evaluate the individual and interactive effects of site, host species, and temperature metrics on symbiont community composition (beta diversity), we performed permutational multivariate analysis of variance (PERMANOVA; Anderson 2017) using the ‘adonis2’ function with 999 permutations (Table S5).

While NMDS, PERMDISP, and PERMANOVA assessed community-level differences, differential abundance testing was conducted to identify specific ASVs that differed significantly between the metrics of interest. The “DESeq2” package was used to assess differential abundance of ASVs across tested factors. A negative binomial generalised linear model (GLM) was fit to raw count data using the ‘DESeq’ function with the parameter ‘fitType’ set to “local” (Fig. S6). Log2 fold changes were calculated for all pairwise group comparisons. Wald tests were then used to assess the significance of differences in symbiont ASV abundance, with Benjamini-Hochberg adjusted p-values set to a 5% false discovery rate. Results were plotted to visualise Symbiodiniaceae taxa driving differences in community composition.

Alpha diversity was calculated using untransformed count data using Shannon’s diversity index (*H’*) and Simpson’s diversity index (1 – *D*). Species richness and evenness of Symbiodiniaceae communities were assessed across categorical factors (host species, sites, and number of accumulated heat stress events; Table S6). Prior to statistical comparisons, data were tested for normality (Shapiro-Wilk Test; Fig. S7, Table S7) and homoscedasticity (Levene’s Test; Table S8) using the “car” package (version 3.1.2; Fox and Weisberg 2019). Depending on data distribution, differences in alpha diversity among groups were evaluated using independent *t*-tests or Welch’s ANOVA (Tables S9, S10).

## Results

### 1. Temperature metrics vary across sites

The reef sites where corals were sampled exhibited distinct patterns across different sea surface temperature (SST) metrics, the number of accumulated heat stress events, and recovery time post heatwave (Fig. 1, Table 1). Maximum Monthly Mean (MMM) temperatures followed a latitudinal gradient, with the southernmost site (Coral Bay = COR) exhibiting the lowest mean temperature (MMM = 26.78 ± 1.85°C SE), whilst the northernmost site (Muiron Islands = MUI) was the warmest (MMM = 28.02 ± 2.04°C; Fig. S8a, File S1). The site located within the Exmouth Gulf (Bundegi = BUN) exhibited the largest temperature range (± 5.77 ± 0.62°C) compared to the six other sites. For the six sites outside of Exmouth Gulf, mean temperature range decreased with latitude (range: 4.95 ± 0.53°C - 5.30 ± 0.59°C; Fig. S8b, File S1). Variability in temperature during the warmest part of the year was lowest at central sites (Osprey Sanctuary = OSP, Lakeside/Turquoise Bay = LK/TR, Tantabiddi = TAN; < 0.57 ± 0.13°C), compared to greater variability during the warmest part of the year at the northern and southern extremes of the sampling range (MUI, VLF, BUN, COR; 0.63 - 0.68 ± 0.13°C; Fig. S8c, File S1). The history of heat stress also varied spatially, with the southernmost site (COR) experiencing the greatest number of accumulated heat stress events (n = 6–9 events; defined as DHW ≥ 8°C-weeks) and the shortest recovery time between events (mean 5.15 ± 0.35 years; File S1). Northern sites (MUI, VLF, BUN) experienced an intermediate number of accumulated heat stress events (n = 3-6 events) and recovery time ranged from 6.98 ± 0.60 years to 13.95 ± 1.21 years. Sites with the lowest number of accumulated heat stress events included the central sites (OSP, LK/TR, TAN; n = 1–3 events), which also recorded recovery times exceeding 12 years (Table 1, File S1).

**Fig. 1.**
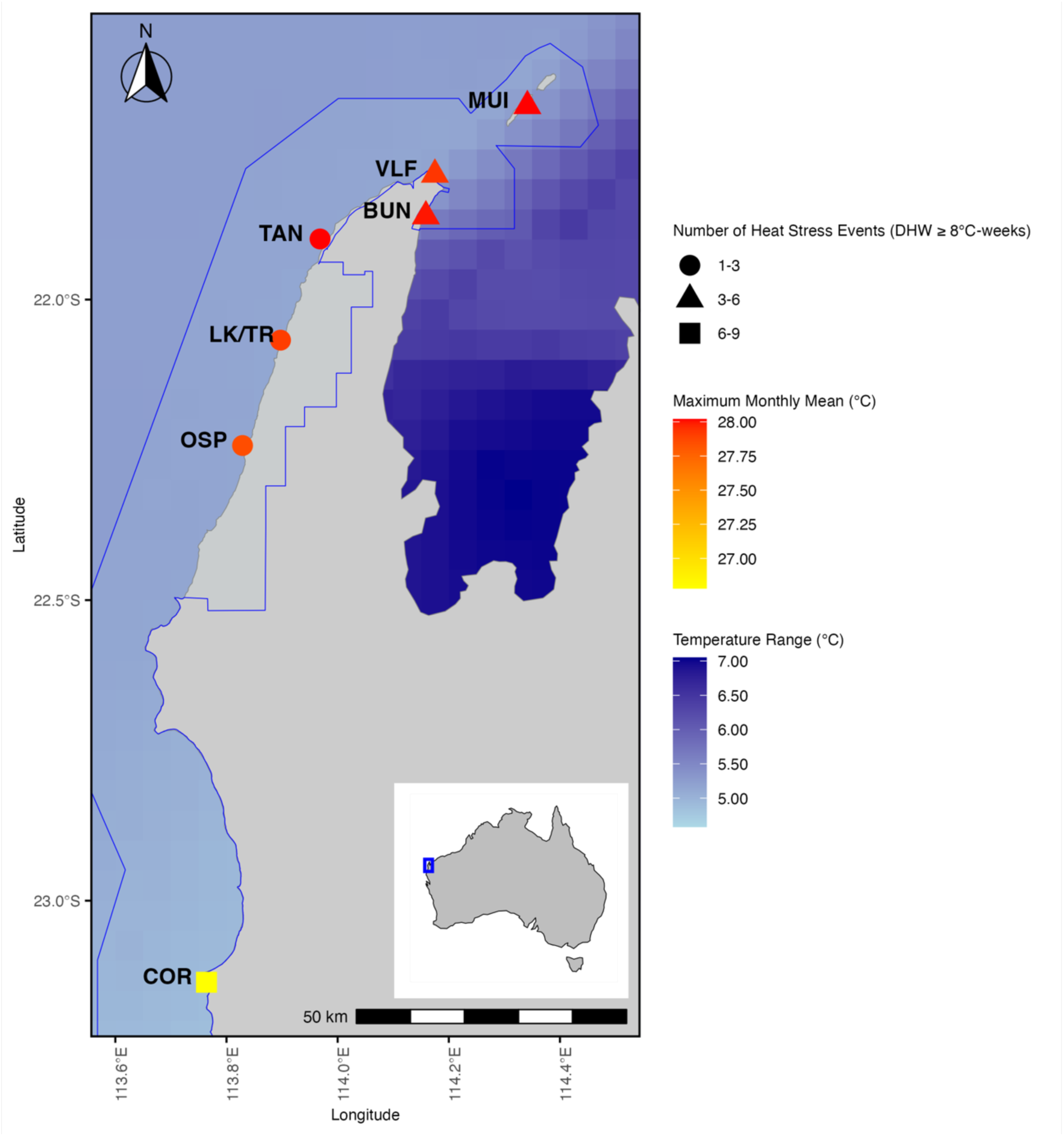
Spatial variation in key temperature metrics across sampled sites along World Heritage listed Ningaloo Reef, Western Australia. The temperature metrics depicted here include Maximum Monthly Mean (MMM) of sea surface temperatures (°C; cooler yellow to warmer red), annual temperature range (°C; less to greater in dark blue) and the number of accumulated heat stress events DHW ≥ 8°C-weeks per site. Shapes correspond to the following: 1–3 events (circles), 3–6 events (triangles) and 6–9 events (squares). Blue outlines indicate UNESCO World Heritage Ningaloo Coast boundaries (UNESCO 2023). The inset map highlights the location of the Ningaloo Reef within Australia

### 2. Symbiodiniaceae community composition and correlation with temperature metrics

Acknowledging the complexity of Symbiodiniaceae taxonomy (see Davies et al. 2023 for a full discussion), groups of similar Amplicon Sequence Variants (ASVs) are described herein as “taxa” for simplicity, though they may not represent formally recognised taxonomic units. Approximately 1,001 ASVs were mapped to 25 taxa within the genus *Cladocopium* across all coral samples. Overall, Symbiodiniaceae communities were dominated by C15, which accounted for 708 of the total ASVs. In terms of normalised relative abundances, this was followed by C116 (49 ASVs), C15h (35 ASVs), and C15.9 (29 ASVs) (Fig. S9), with the remaining 180 ASVs belonging to other *Cladocopium* taxa.

The prevalence of different *Cladocopium* taxa and their normalised relative abundance varied across sites (Fig. 2, File S2). Specific taxa were unique to certain sites. For example, C15.1 was only detected in BUN, C91a in COR, C15j and C50 in MUI, while other taxa were absent in other sites (e.g. C116 and C15.8 were not present in LK/TR and OSP, or BUN, respectively). Other *Cladocopium* taxa were ubiquitous across all sites and host corals (Fig. 2, Table S11). C15 was detected across all samples and sites, comprising 89 ± 1.6% of the relative abundance of ASVs per sample (Tables S3, S12).

**Fig. 2.**
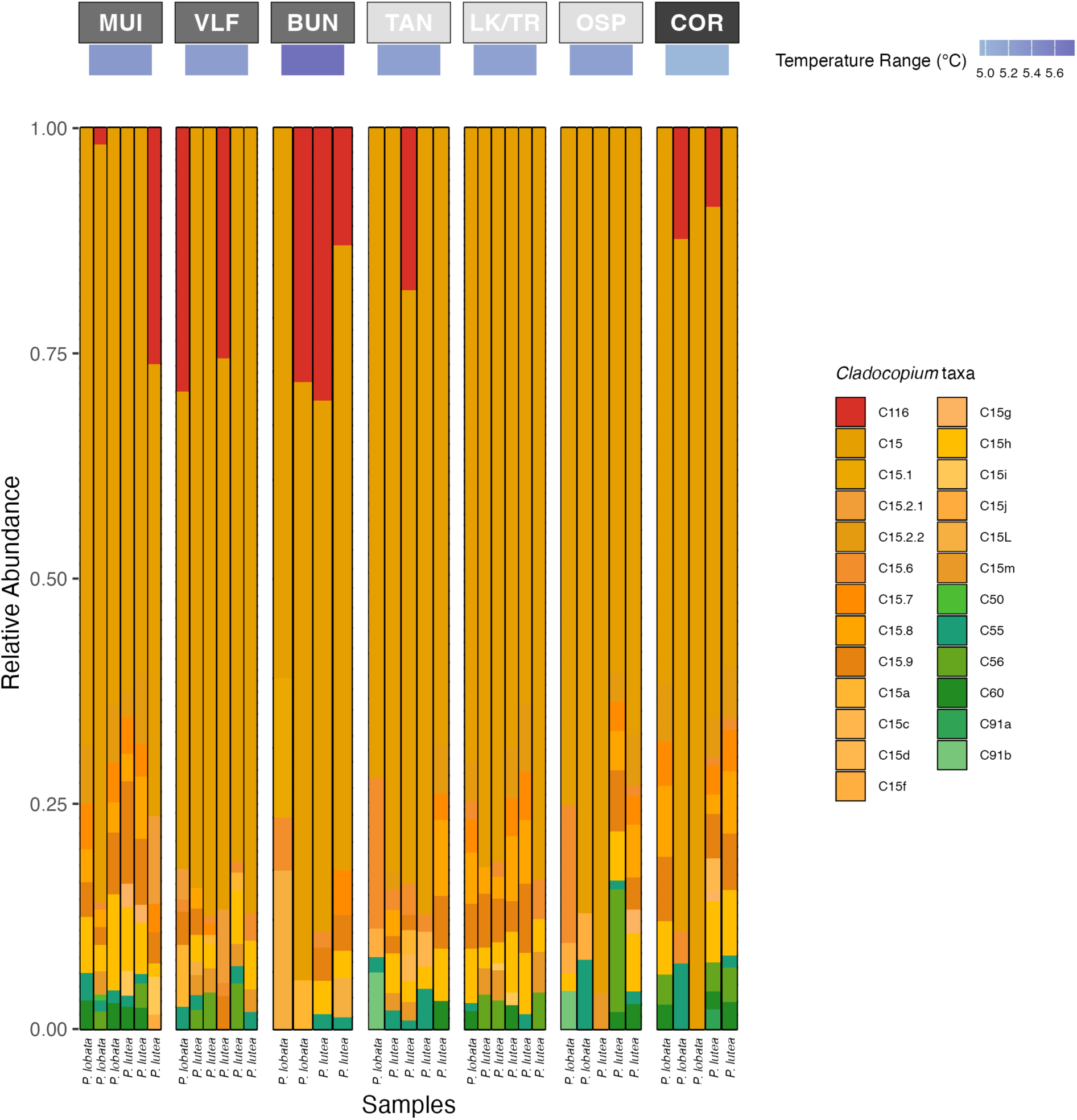
Normalised relative abundances of *Cladocopium* taxa across samples and sites. Each bar represents a *Porites* coral sample. Different colours within each bar represent the *Cladocopium* taxa detected in each coral sample. Each bar is labelled by host species (bottom). Site labels (top) are arranged from north to south (from left to right), with the Muiron Islands (MUI) as the northernmost site. Site labels, in grey-scale, correspond to the number of accumulated heat stress events DHW ≥ 8°C-weeks per site (1–3 events (light grey), 3–6 events (grey) and 6–9 events (dark grey). The colour beneath each site label indicates the annual temperature range experienced at that site (temperature range °C; lower temperatures in light purple to higher temperatures dark purple)

C15h, the second most prevalent *Cladocopium* taxon, was detected in 28 of 37 samples, with a mean relative abundance of 0.7 ± 0.1% per sample (Table S12). C15h was detected across all sites and was most abundant at the warmest site (highest MMM, MUI, Fig. 2, File S2). At the site with the greatest temperature range (BUN), C15h was less common and only found in 50% of samples. C15h was more common in *P*. *lutea* compared to *P*. *lobata* (found in 92% compared to 46% of samples; File S2). Another common taxon, C15.9, was found in over 40% of samples per site (File S2), with a mean relative abundance of 1.4 ± 0.2% (Table S12). It was most prevalent at the warmest site and was more common in *P*. *lutea* compared to *P*. *lobata* (75% vs. 46%, File S2).

Of the Symbiodiniaceae taxa, 28% (7 of 25) of the *Cladocopium* were not identified to C15. Six of seven were in low abundance (≤ 1%, Table S12). C116 was the exception, with this taxon present in 10 of the total 37 samples across five sites (Tables S3, File S2), and a mean relative abundance of 12.6 ± 3.0% across samples (Table S12). C116 was most common at the site with the largest temperature range (BUN), where it made up 13–30% of the relative abundance in 75% of samples (Fig. 2). In the site with the smallest temperature range and greatest number of accumulated heat stress events (COR), C116 accounted for >10% of the relative abundance in 40% of samples. At sites that experienced an intermediate to high number of accumulated heat stress events (3-6 and 6-9 events, respectively), C116 was found in both host species (Fig. 2). C116 was uncommon or absent at sites that experienced the lowest number of accumulated heat stress events (1-3 events; Fig. 2, TAN, LK/TR, OSP).

Several low-abundance *Cladocopium* taxa were broadly distributed across sites and host species. C55 was present in 22 of 37 samples, with a mean relative abundance of 0.3 ± 0.1%, occurring in 62% of *P*. *lobata* and 58% of *P*. *lutea* samples (Table S12, File S2). It was most frequent at the warmest sites of highest MMM (MUI and TAN, ∼80% of samples) and commonly co-occurred with C15.9 and/or C15h (19 samples; Table S3). C56 (1.3 ± 0.6%) and C60 (0.3 ± 0.04%) were detected in 12 samples across five sites (Tables S3, S12) and were more common in *P*. *lutea* (42% and 33%, respectively) than *P*. *lobata* (15% and 31%, respectively). Both C56 and C60 were more prevalent (60% of samples) at the site with the smallest temperature range and the greatest number of accumulated heat stress events (COR). Notably, neither C56 nor C60 were detected at the site with the largest temperature range (BUN; Table S11).

Overall, *Cladocopium* taxa exhibited strong spatial structuring and were significantly correlated with these subsets of metrics, including temperature range, latitude, and number of accumulated heat stress events (Figs. 1, 2). Variation in Symbiodiniaceae community composition was significantly explained by site-specific differences in temperature metrics, as shown through non-metric multidimensional scaling (NMDS, Fig. 3a). Specifically, 24% of variation in community composition was explained by site (PERMANOVA: *F* = 1.586, *R²* = 0.2408, *p* = 0.004; Table S5). Of the temperature metrics, the number of accumulated heat stress events explained the greatest variation in symbiont community structure (PERMANOVA: *F* = 1.604, *R²* = 0.0862, *p* = 0.041; Fig. S10a). This was followed by temperature range (PERMANOVA: *F* = 2.436, *R²* = 0.0651, *p* = 0.002; Figs. 3b, S10b) and a small but significant proportion of variation in symbiont community structure was explained by latitude (< 5%, PERMANOVA: *F* = 1.792, *R²* = 0.0487, *p* = 0.036; Fig. S10c). The remaining metrics (MMM, variation in the warmest time of year, recovery time, or longitude) did not significantly explain differences in symbiont community composition (PERMANOVA: *p >* 0.05; Table S5).

**Fig. 3.**
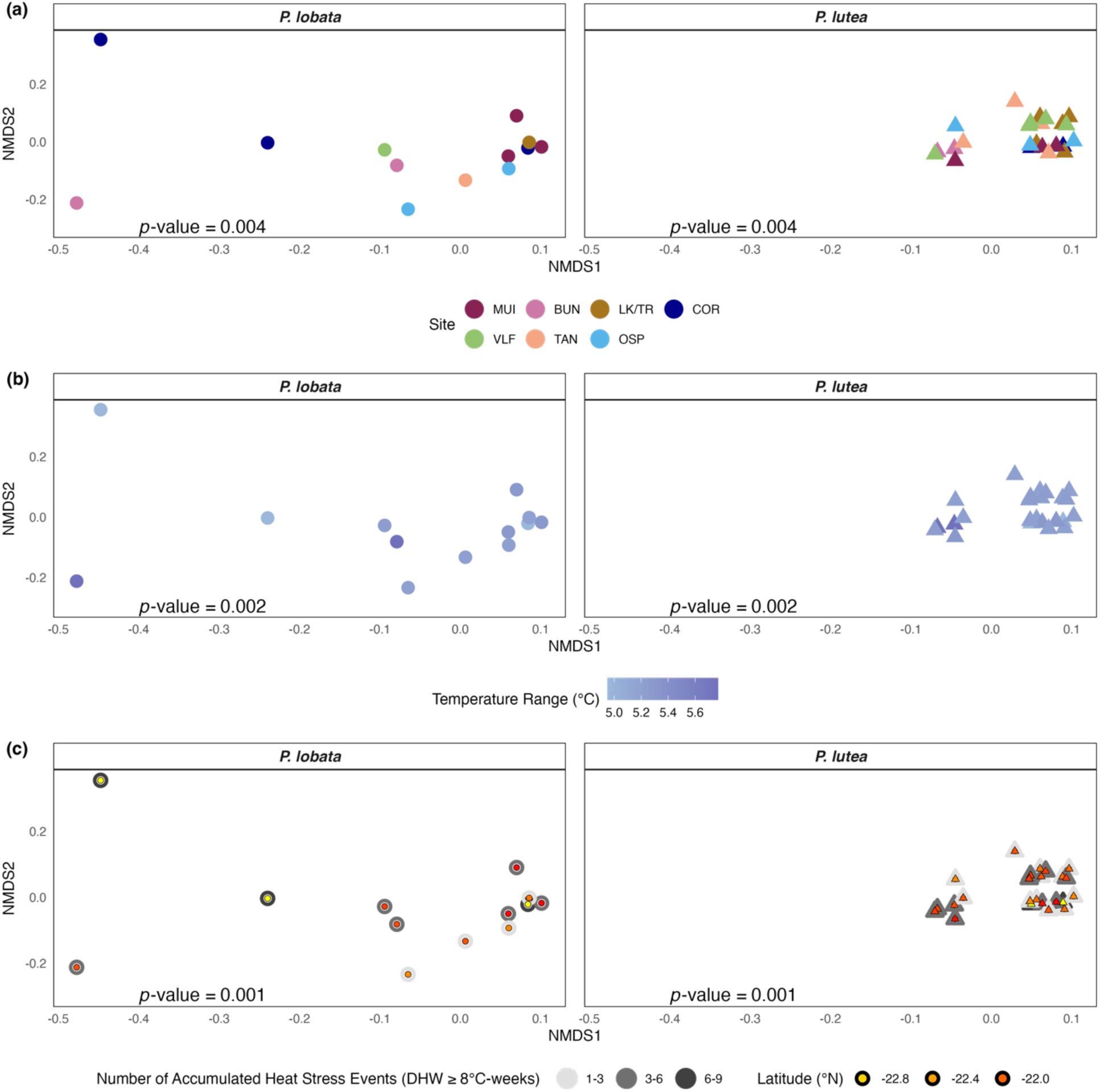
Non-metric multidimensional scaling (nMDS) of Symbiodiniaceae community composition based on weighted UniFrac distances. Ordinations display Symbiodiniaceae community composition for *P*. *lobata* (circles) and *P*. *lutea* (triangles) across environmental factors. Community dispersion and structure were significantly different between host species (Homogeneity of Dispersions: *F* = 13.113, *p* = 0.001; PERMANOVA, *p* = 0.007). Communities are coloured by **(a)** site (PERMANOVA, *p* = 0.004) **(b)** temperature range (°C; PERMANOVA, *p* = 0.002), and **(c)** number of heat stress events x latitude (DHW ≥ 8°C-weeks x °N; PERMANOVA, *p* = 0.001)

The interaction between selected temperature metrics significantly explained variation in symbiont community structure across sites (Table S5). Significant interactions included temperature range × warm season variability (PERMANOVA: *F* = 1.822, *R²* = 0.04756, *p* = 0.034) and temperature range × recovery time (PERMANOVA: *F* = 1.911, *R²* = 0.0497, *p* = 0.029). Notably, interactions between the number of accumulated heat stress events and latitude (PERMANOVA: *F* = 2.780, *R²* = 0.1354, *p* = 0.001), as well as longitude (PERMANOVA: *F* = 2.445, *R²* = 0.1204, *p* = 0.001), collectively explained over 25% of variation in community structure, and were reflected in patterns of clusters across ordination space (Fig. 3c).

Symbiodiniaceae relative abundance varied across sites depending on the number of accumulated heat stress events (Figs. 2, 4). Specifically, ASVs mapping to 10 *Cladocopium* taxa were significantly differentially abundant across pairwise comparisons between sites experiencing 1-3, 3-6, and 6-9 accumulated heat stress events (Log₂ fold change: padj < 0.05; Fig. 4, File S2). Notably, three of these taxa were recorded exclusively within specific heat stress categories (Table S13). For example, C15.2.1 was only detected at sites with an intermediate number of events (MUI, VLF; n = 3-6 events) and C15f was detected only at sites with the lowest number of events (TAN, OSP; n = 1-3 events). ASVs identified as C15.2.1 and C15f were significantly more abundant in these respective categories (Fig. 4). C15m was present at sites with low to intermediate heat stress (1-3 and 3-6 events) and showed significantly higher abundance in both 1-3 and 3-6 events compared to 6-9 events. However, its abundance did not differ significantly between 1-3 and 3-6 event categories (Fig. 4). Seven of these 10 *Cladocopium* taxa were observed across all three categories (Table S13). ASVs belonging to C56 decreased in abundance proportionally with the number of events (1–3: 2.9%, 3–6: 2.2%, 6–9: 2%; Table S13), which was reflected in pairwise comparisons between categories (Log_2_ fold change: *p.adj <* 0.05; Fig. 4). Alternatively, ASVs belonging to C116 increased proportionally with the number of events (1–3: 2.4%, 3–6: 6.5%, 6–9: 7.0%, Table S13). These C116 ASVs were significantly more abundant with an intermediate number of accumulated heat stress events (Fig.4).

**Fig. 4.**
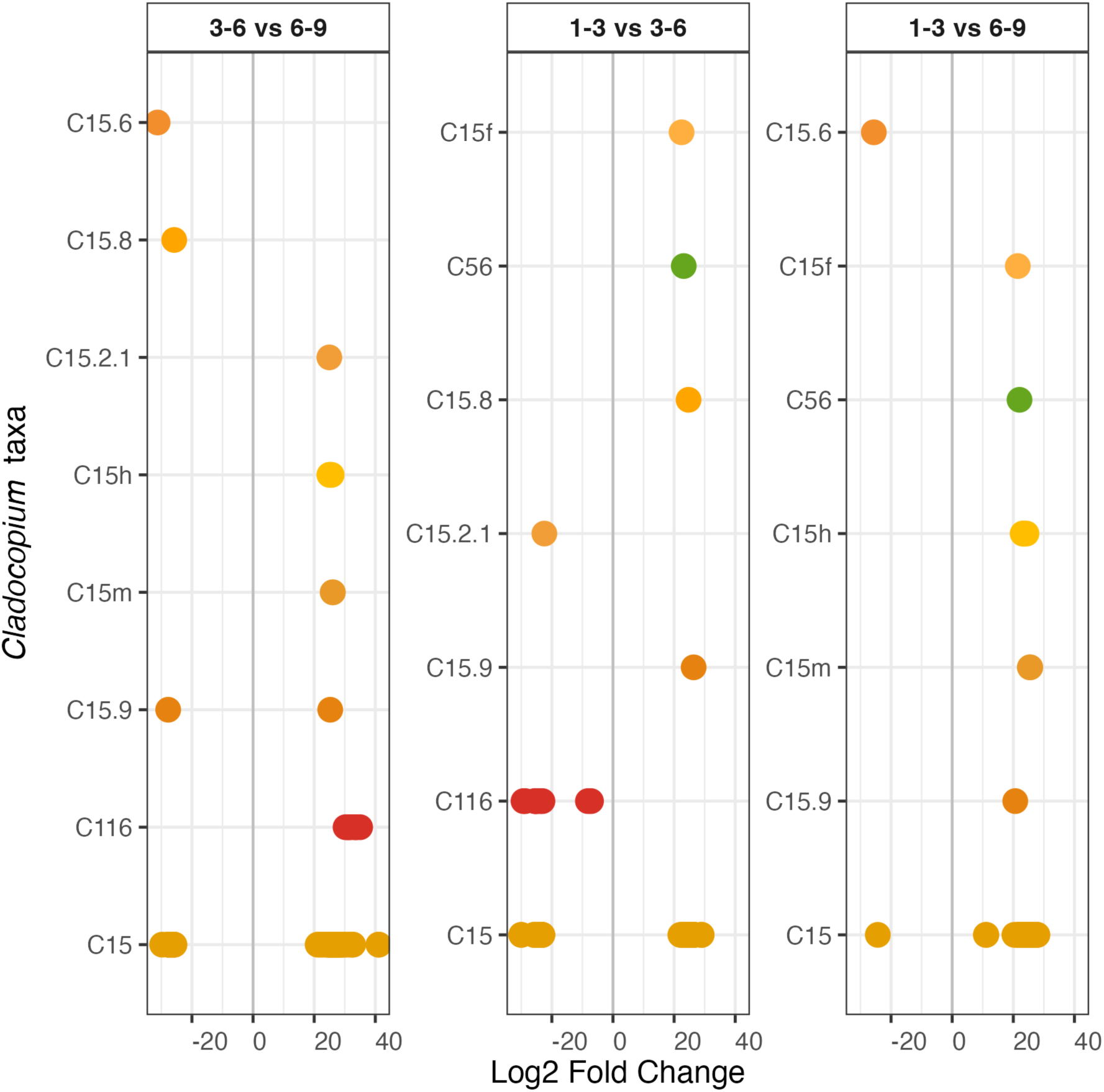
Change in abundance of specific *Cladocopium* taxa across categories of the number of accumulated heat stress events DHW ≥ 8°C-weeks. Each panel represents a pairwise comparison of 1–3 vs. 6–9 events, 3–6 vs. 6–9 events, and 1–3 vs. 3–6 events. Each of the taxa represented here were significantly differentially abundant (adjusted p-value < 0.05), represented as log2 fold-change between categories. Positive fold changes indicate higher abundance in the first category compared to the second, while negative values indicate higher abundance in the second category. Multiple points presented per *Cladocopium* taxa represent multiple significantly differentially abundant ASVs identified within each taxon

### 3. Symbiodiniaceae communities differ by host species

Variation in Symbiodiniaceae community composition varied significantly by host species (PERMDISP: *F* = 13.113, *p* = 0.001; PERMANOVA: *F* = 2.181, *R²* = 0.0587, *p* = 0.007; Tables S4, S5). Specific *Cladocopium* taxa were uniquely associated with each host species. For example, C15f and C91b were only detected in *P*. *lobata* samples, whereas C15g, C15i, C15d were only detected in *P*. *lutea* samples (Table S11). Seven of the 15 *Cladocopium* taxa detected in both host species (e.g., C15, C55, C60, C15.6, C15.2.1) were observed at similar frequencies across *P*. *lutea* and *P*. *lobata* samples. The other eight shared taxa had a higher prevalence in *P*. *lutea* samples (File S2). However, communities in *P*. *lutea* were significantly less variable compared to *P*. *lobata*, as was reflected in cluster patterns across ordination space (Fig. 3).

### 4. Sequence diversity within Symbiodiniaceae communities

To further explore sequence-based diversity, two alpha diversity metrics were calculated using different factors (host species, site, and number of accumulated heat stress events). These metrics included Shannon’s Diversity Index (hereafter referred to as richness; *H’*) and Inverted Simpson’s Index (hereafter referred to as evenness; 1-*D*). Richness was greater in *P*. *lutea* compared to *P*. *lobata* (*H’* = 2.272 ± 0. 09 vs. 2.121 ± 0.14), whereas evenness was greater in *P*. *lobata* compared to *P*. *lutea* (1-*D* = 0.775 ± 0.03 vs. 0.772 ± 0.02; Fig. S11a, Table S6). These differences were not significant (independent t-test, *p >* 0.05; Table S9). Trends in these metrics across sites were present but not statistically significant (Welch’s ANOVA, *p* > 0.05; Table S10).

Richness was lowest at the coolest sites (COR, *H’* = 2.052 ± 0.19) and highest at the sites of highest MMM (MUI, *H’* = 2.443 ± 0.17; Fig. S11b). Evenness was lowest at the most central site (LK/TR, 1-*D* = 0.747 ± 0.03) and highest at the warmest site (MUI, 1-*D* = 0.802 ± 0.03; Fig. S11b). Sites with an intermediate number of accumulated heat stress events (3-6 events) exhibited the highest richness and evenness (*H’* = 2.285 ± 0.14 and 1-*D* = 0.785 ± 0.03; Fig. S11c). Sites with the highest number of accumulated heat stress events (6–9 events) had the lowest richness (*H’* = 2.052 ± 0.19) and sites with the lowest number of accumulated heat stress events (1-3 events) had the lowest evenness (1-*D* = 0.762 ± 0.02; Fig.S11c).

## Discussion

This study provides novel insights into the diversity and distribution of Symbiodiniaceae communities in massive *Porites* species (*P*. *lutea* and *P*. *lobata*) along the Ningaloo World Heritage reef in Western Australia. We found that symbiont community composition varied both by site and host species, with patterns closely associated with sea surface temperature (SST) variation and the frequency of accumulated heat stress events (DHW ≥ 8°C-weeks). Sites with greater temperature fluctuations and more frequent thermal stress (such as Bundegi and Coral Bay) harboured higher abundances of thermotolerant *Cladocopium* taxa, notably C116.

These findings suggest a potential shift in symbiont assemblages towards heat-tolerant types in more impacted areas. The presence of C116 at warmer, or more disturbed, sites may reflect prior exposure to stress (Swain et al. 2016). In contrast, at historically thermally stable sites (Lakeside/Turquoise Bay, Osprey Sanctuary), C116 was either present in low abundance or absent altogether, suggesting it may only become abundant in response to thermal stress. This pattern suggests that C116 may play a role in enhancing coral resilience in regions exposed to recurrent heat stress.

Interestingly, while *Cladocopium* C15 was the dominant symbiont across most samples and sites, we also recovered substantial diversity among C15 associated ASVs, each present at varying abundances. Although Symbiodiniaceae taxonomy is under active revision (Davies et al. 2023), this diversity may reflect the presence of distinct lineages within the broader C15 group. Further support for this hypothesis comes from the detection of these ASVs across different heat stress categories. If these ASVs represent different lineages, their distribution may reflect local adaptation to varying temperature conditions observed across our sample sites, as has been observed in other studies around the world (Arif et al. 2014; Claar et al. 2020). Moreover, aside from the temperature-related metrics examined here, other factors such as connectivity across sites, distance from shore, exposure to the Leeuwin Current or tidal currents may also play a role (Vanderklift et al. 2020) in influencing symbiont community structure, aligning with previous findings for massive *Porites* (Tan et al. 2020).

Notably, a range of background taxa, including C50 and C91a were also detected. This indicates that *Porites* species are not only dominated by C15 but may also benefit from the additional functional diversity contributed by other Symbiodiniaceae taxa. The trend of detecting additional diversity within *Porites* species is growing with improved sequencing tools (see Claar et al. 2020). While the ecological roles of these background taxa are not fully understood, they may play important roles in providing functional redundancy and enhancing holobiont stability under differing environmental conditions (Ziegler et al. 2018).

We also found that symbiont community variability was significantly correlated with host species. While *P*. *lobata* communities exhibited greater variability across sites, particularly at sites experiencing greater heat stress and a broader temperature range, *P*. *lutea* harboured a greater abundance of diverse symbiont taxa within individual colonies. Differences in symbiont community structure between these two host species may reflect differences between *P*. *lutea* and *P*. *lobata* in terms of their ecological niches or strategies for coping with environmental stress (Grupstra et al. 2024). These preliminary results suggest that *P*. *lobata* may rely on a broader range of symbiont associations across their geographic range to respond to their environment, whereas *P*. *lutea* may maintain a higher abundance of specific taxa to buffer against environmental stress. Moreover, these differences are likely driven in part by host genetics (Starko et al. 2023; Grupstra et al. 2024) or are an emergent property of differences in physiological traits like tissue thickness (Putnam et al. 2017). The pervasiveness of both shared and unique C15 variants across these two host species further supports the hypothesis that different *Cladocopium* taxa likely fulfil distinct functional roles.

These species-specific findings help to support challenges against the long-held paradigm that *Porites* species host only stable, low-diversity symbiont communities dominated by *Cladocopium* C15 (Putnam et al. 2012). Instead, our results here and others (Qin et al. 2019) show that *Porites* species exhibit greater symbiont flexibility and diversity than previously assumed (Terraneo et al. 2024). This may be contributing to their resilience to thermal stress. Indeed, the ability to adjust the relative abundance of existing symbionts (“shuffling”) or to acquire new symbiont types (“switching”) are common mechanisms corals use to cope with environmental stress (Baker 2003; Quigley et al. 2022). The detection of thermally tolerant taxa like *Cladocopium* C116 suggests that *Porites* may have greater capacity than initially thought to shuffle or switch in response to heat stress or differences across the environment (Starko et al. 2023). However, we also observed declines in key symbiont diversity metrics, including richness and evenness at both high heat stress sites (Coral Bay) and more thermally stable sites (Lakeside). These findings may indicate limits to this flexibility or resilience, but further research is needed to confirm the causes and consequences of these declines in diversity metrics.

Given the increasing frequency and severity of marine heat waves globally, it is critical to understand how coral-symbiont associations shift across space and time. Our study highlights previously undetected diversity and flexibility in Symbiodiniaceae communities hosted by massive *Porites* species along the Ningaloo Coast. Importantly, we found that heat stress history and temperature variability may play a key role in driving the structure of these symbiont communities, with likely long-term impacts on reef resilience. Continued monitoring of these associations is essential for predicting how reef-building corals and their symbiotic partners will respond under future climate regimes. Future research should explore the functional roles of background and thermotolerant symbiont taxa across broader spatial and temporal scales to assess their contributions to holobiont performance and long-term reef persistence in the eastern Indian Ocean and beyond.

## Statements and Declarations

### Funding

This research was conducted under the Minderoo Foundation grants: GCA Porites - 346626.4 to Z.T.R. and D.J., Putting MERL on the Map to K.M.Q and was a collaborative effort between James Cook University, Minderoo Foundation, Curtin University, and the Department of Biodiversity, Conservation, and Attractions. K.M.Q. is supported by funding from ARC DECRA Fellowship (DE230100284) and Minderoo Foundation.

### Ethical Approval

Samples were collected under DBCA Fauna Taking Licenses #FO25000342-3 and #FO25000458 and Department of Primary Industries and Regional Development, Western Australia Exemptions #251145223 and #251143023.

### Conflict of Interest

The authors declare no conflicts of interest.

### Contributions

All authors contributed to the conception and design of the study. K.M.Q., D.J., and Z.T.R. obtained permits, permissions, and funding for the project. L.K. and D.J. conducted fieldwork and lab processing of samples. D.J. performed *Porites* morphological assessment and host species identification. L.K. performed DNA extraction, quantification, PCR amplification, and analysis of Symbiodiniaceae data. L.K. wrote the original draft of the manuscript and all authors reviewed and provided edits to the manuscript. All authors approved the final version of the manuscript for publication.

### Data and Code Availability Statement

Datasets and scripts that support the results of this study can be accessed through Github [https://github.com/lindseykraemer/Ningaloo_Porites]. Sequencing data will be made available at the time of publication.

## Acknowledgements

We respectfully acknowledge the Baiyungu, Thalanyji, Yinigurdira, Bindal, Noongar Wadjak and Wulgurukaba people, the Traditional Owners and custodians of the land and sea country upon which our research was conducted. We are grateful to the present and former Minderoo Exmouth Research Laboratory staff members and our dedicated volunteers for their involvement in field operations. We extend a special thanks to the Conservation ‘Omics lab, Mahsa Mousavi-Derazmahalleh, and Niko Andreakis for the lab and bioinformatic support.

## Notes

### Competing Interest Statement

The authors have declared no competing interest.

