## Supplemental Tables (S1-S13), Figures (S1-S11), Data (S1, S2) for "Past thermal history and heat stress shape patterns of coral-algal symbioses in massive *Porites*"

**Supplementary Tables**

**Table S1. Sample and read count information through “DADA2” amplicon analysis workflow.**

| **Sample Name** | **Morpho. Host ID** | **PCR Dilution** | **Input** | **Filtered** | **Denoised** | **Merged** | **Tabled** | **Nonchim** |
| --- | --- | --- | --- | --- | --- | --- | --- | --- |
| NING.BUND.POR.1 | P. cf. lobata | NA | 141414 | 129337 | 129215 | 125392 | 125151 | 117926 |
| NING.BUND.POR.2 | P. lobata | NA | 236237 | 224469 | 224271 | 221155 | 221017 | 207341 |
| NING.BUND.POR.4 | P. lutea | NA | 172423 | 163085 | 162926 | 160882 | 160425 | 152383 |
| NING.BUND.POR.6 | P. lutea | NA | 212090 | 199655 | 199281 | 195831 | 195416 | 173365 |
| NING.BUND.POR.9 | P. australiensis | NA | 203241 | 192619 | 192169 | 188184 | 187702 | 180214 |
| NING.BUND.POR.10 | P. lobata | NA | 117608 | 111088 | 111079 | 110740 | 110476 | 110476 |
| NING.VLF.POR.1 | P. lutea | NA | 244627 | 234020 | 233810 | 225706 | 224904 | 210234 |
| NING.VLF.POR.7 | P. lobata | NA | 243519 | 230219 | 230168 | 224180 | 223330 | 213593 |
| NING.VLF.POR.12 | P. lutea | NA | 226833 | 216303 | 215863 | 209685 | 208559 | 199036 |
| NING.VLF.POR.13 | P. lutea | NA | 222128 | 210479 | 210434 | 205442 | 204884 | 192191 |
| NING.VLF.POR.15 | P. lutea | NA | 214812 | 203857 | 203359 | 196925 | 195725 | 181167 |
| NING.VLF.POR.17 | P. lutea | NA | 220644 | 209250 | 209119 | 206871 | 206774 | 198736 |
| NING.OSP.POR.2 | P. lutea | 1:100 | 182917 | 172573 | 172316 | 166176 | 165443 | 160123 |
| NING.OSP.POR.3 | P. cf. solida | 1:10 | 184737 | 174613 | 174165 | 166777 | 166237 | 159433 |
| NING.OSP.POR.9 | P. lutea | 1:100 | 190599 | 176680 | 176292 | 169085 | 168648 | 165849 |
| NING.OSP.POR.11 | P. lobata | 1:10 | 194482 | 184237 | 183951 | 177118 | 176733 | 164964 |
| NING.OSP.POR.13 | P. lutea | 1:10 | 196571 | 185291 | 185071 | 179254 | 178820 | 169603 |
| NING.OSP.POR.15 | P. lobata | 1:10 | 255811 | 239600 | 239486 | 234759 | 230733 | 219855 |
| NING.TAN.POR.1 | P. lutea | NA | 268696 | 253027 | 252993 | 244100 | 243291 | 225334 |
| NING.TAN.POR.4 | P. lobata | NA | 221935 | 211482 | 211114 | 204679 | 203974 | 190701 |
| NING.TAN.POR.10 | P. lutea | NA | 200034 | 191160 | 190965 | 185663 | 185089 | 169971 |
| NING.TAN.POR.11 | P. lutea | NA | 268872 | 257006 | 256453 | 248051 | 247117 | 240618 |
| NING.TAN.POR.15 | P. australiensis | NA | 244345 | 232704 | 232498 | 226471 | 225721 | 208010 |
| NING.TAN.POR.17 | P. lutea | NA | 201092 | 188385 | 188167 | 184641 | 184414 | 148773 |
| NING.MUI.POR.4 | P. lutea | NA | 227734 | 214309 | 214294 | 171650 | 170822 | 169498 |
| NING.MUI.POR.10 | P. lutea | 1:10 | 287438 | 274429 | 274389 | 272119 | 271606 | 267804 |
| NING.MUI.POR.14 | P. lutea | 1:10 | 246903 | 235028 | 234848 | 188657 | 187773 | 185777 |
| NING.MUI.POR.17 | P. lobata | NA | 274456 | 261868 | 261542 | 253310 | 250681 | 241111 |
| NING.MUI.POR.18 | P. lobata | NA | 253457 | 236423 | 236351 | 176682 | 175289 | 174735 |
| NING.MUI.POR.20 | P. lobata | 1:10 | 203662 | 194202 | 194108 | 187727 | 185546 | 177571 |
| NING.COR.POR.5 | P. lobata | 1:10 | 216089 | 204732 | 204327 | 196652 | 195687 | 186107 |
| NING.COR.POR.8 | P. cf. lutea | NA | 193521 | 184433 | 183909 | 179938 | 179640 | 169897 |
| NING.COR.POR.9 | P. lobata | NA | 225870 | 208914 | 208561 | 201107 | 200432 | 191606 |
| NING.COR.POR.13 | P. lutea | 1:10 | 190168 | 179761 | 179522 | 176560 | 176043 | 165767 |
| NING.COR.POR.18 | P. lutea | 1:100 | 199733 | 190450 | 190206 | 184763 | 184442 | 170482 |
| NING.COR.POR.20 | P. lobata | 1:10 | 236371 | 220326 | 220195 | 213437 | 212472 | 199042 |
| NING.TUR.POR.1 | P. lutea | NA | 197550 | 188662 | 187856 | 178536 | 176223 | 161517 |
| NING.TUR.POR.2 | P. lobata | NA | 241872 | 229804 | 229601 | 223977 | 223214 | 207040 |
| NING.LAK.POR.1 | P. lutea | NA | 184359 | 174103 | 174008 | 166853 | 165968 | 161108 |
| NING.LAK.POR.2 | P. lutea | NA | 187415 | 177757 | 177632 | 173173 | 172866 | 169936 |
| NING.LAK.POR.3 | P. cf. lobata | 1:10 | 196328 | 185895 | 185552 | 182290 | 180833 | 161917 |
| NING.LAK.POR.4 | P. lutea | 1:10 | 231878 | 218159 | 217777 | 213492 | 212194 | 207084 |
| NING.LAK.POR.5 | P. lutea | 1:10 | 204022 | 192735 | 192317 | 183876 | 182393 | 171663 |
| NING.LAK.POR.6 | P. cf. lutea | 1:10 | 186335 | 176748 | 176519 | 173235 | 173037 | 165456 |

*Note:* For each sequenced sample (Sample Name; n = 44), columns from left to right represent morphologic identification (Morpho. Host ID), final PCR dilution (PCR Dilution), and read count after sequential “DADA2” steps, as follows: read yield after trimming residual adapter and index content (Input), filtered reads (Filtered), inferred sequence variants (Denoised), merged pair reads (Merged), ASVs that fell within expected length range (294-304 bp) (Tabled), non-chimeric sequences (Nonchim).

**Table S2. BLAST Results for ‘NA’ ASV Sequences.**

| **ASV ID** | **Genbank ID** | **Percent Identity** | **E value** | **Accession No.** |
| --- | --- | --- | --- | --- |
| seq10 | C15 | 99.67 | 2.00E-151 | JN558044.1 |
| seq14 | C15 | 99.33 | 8.00E-150 | JN558044.1 |
| seq20 | C15 | 99.34 | 8.00E-150 | JN558044.1 |
| seq38 | C3 | 98.33 | 4.00E-145 | MN265288.1 |
| seq52 | C15 | 99.67 | 2.00E-151 | JN558044.1 |
| seq56 | C15 | 99.67 | 2.00E-151 | JN558044.1 |
| seq65 | C15 | 99.67 | 2.00E-151 | JN558044.1 |
| seq72 | C15 | 99.67 | 2.00E-151 | JN558044.1 |
| seq76 | C15 | 98.67 | 2.00E-146 | JN558044.1 |
| seq80 | C15 | 99.67 | 2.00E-151 | JN558044.1 |
| seq81 | C15 | 99.67 | 2.00E-151 | JN558044.1 |
| seq82 | C15 | 99.67 | 2.00E-151 | JN558044.1 |
| seq87 | C15 | 99.67 | 2.00E-151 | JN558044.1 |
| seq104 | C15 | 99.67 | 2.00E-151 | JN558044.1 |
| seq107 | C15 | 99.33 | 8.00E-150 | JN558044.1 |
| seq112 | C15 | 99.67 | 2.00E-151 | JN558044.1 |
| seq117 | C15 | 99.67 | 2.00E-151 | JN558044.1 |
| seq123 | C15 | 99 | 4.00E-148 | JN558044.1 |
| seq129 | C15 | 99.67 | 2.00E-151 | JN558044.1 |
| seq135 | C15 | 99.67 | 2.00E-151 | JN558044.1 |
| seq139 | C15 | 99.67 | 2.00E-151 | JN558044.1 |
| seq141 | C15 | 100 | 5.00E-152 | JN558044.1 |
| seq186 | C15 | 99.67 | 2.00E-151 | JN558044.1 |
| seq189 | C15 | 99.67 | 2.00E-151 | JN558044.1 |
| seq195 | C15 | 99.67 | 2.00E-151 | JN558044.1 |
| seq203 | C15 | 99.34 | 8.00E-150 | JN558044.1 |
| seq215 | C3 | 99 | 4.00E-148 | MN876157.1 |
| seq216 | C3 | 99 | 4.00E-148 | MN876157.1 |
| seq218 | C15 | 99.33 | 8.00E-150 | JN558044.1 |
| seq221 | C15 | 99.67 | 2.00E-151 | JN558044.1 |
| seq226 | C15 | 99.33 | 8.00E-150 | JN558044.1 |
| seq229 | C15 | 99.67 | 2.00E-151 | JN558044.1 |
| seq261 | C15 | 99.67 | 2.00E-151 | JN558044.1 |
| seq268 | C15 | 99.67 | 2.00E-151 | JN558044.1 |
| seq277 | C15 | 99.33 | 8.00E-150 | JN558044.1 |
| seq283 | C15 | 99.33 | 8.00E-150 | JN558044.1 |
| seq284 | C15 | 98.67 | 2.00E-146 | JN558044.1 |
| seq286 | C15 | 99.67 | 2.00E-151 | JN558044.1 |
| seq293 | C15 | 99.33 | 8.00E-150 | JN558044.1 |
| seq309 | C15 | 99.33 | 8.00E-150 | JN558044.1 |
| seq311 | C15 | 99.67 | 2.00E-151 | JN558044.1 |
| seq327 | C15 | 99.67 | 2.00E-151 | JN558044.1 |
| seq342 | C15 | 99 | 4.00E-148 | JN558044.1 |
| seq365 | C15 | 99.67 | 6.00E-151 | JN558044.1 |
| seq379 | C15 | 99.33 | 8.00E-150 | JN558044.1 |
| seq380 | C15 | 99.67 | 2.00E-151 | JN558044.1 |
| seq389 | C15 | 99.67 | 2.00E-151 | JN558044.1 |
| seq397 | C15 | 99.67 | 2.00E-151 | JN558044.1 |
| seq403 | C15 | 99 | 4.00E-148 | JN558044.1 |
| seq404 | C15 | 99.67 | 6.00E-151 | JN558044.1 |
| seq407 | C15 | 99.33 | 3.00E-149 | JN558044.1 |
| seq412 | C15 | 99 | 4.00E-148 | JN558044.1 |
| seq420 | C15 | 99.67 | 6.00E-151 | JN558044.1 |
| seq430 | C15 | 99.33 | 8.00E-150 | JN558044.1 |
| seq432 | C15 | 99.67 | 6.00E-151 | JN558044.1 |
| seq433 | C15 | 99 | 4.00E-148 | JN558044.1 |
| seq459 | C3 | 100 | 4.00E-153 | MN876157.1 |
| seq464 | C15 | 99 | 1.00E-147 | JN558044.1 |
| seq466 | C15 | 99.33 | 8.00E-150 | JN558044.1 |
| seq469 | C15 | 99.67 | 6.00E-151 | JN558044.1 |
| seq475 | C15 | 99.67 | 2.00E-151 | JN558044.1 |
| seq483 | C15 | 99 | 4.00E-148 | JN558044.1 |
| seq494 | C15 | 99.67 | 2.00E-151 | JN558044.1 |
| seq497 | C15 | 99.67 | 6.00E-151 | JN558044.1 |
| seq503 | C15 | 99.67 | 6.00E-151 | JN558044.1 |
| seq508 | C15 | 99.66 | 8.00E-150 | JN558044.1 |
| seq509 | C15 | 99.33 | 3.00E-149 | JN558044.1 |
| seq510 | C15 | 99.33 | 8.00E-150 | JN558044.1 |
| seq514 | C15 | 99.66 | 2.00E-150 | JN558044.1 |
| seq515 | C15 | 99.67 | 6.00E-151 | JN558044.1 |
| seq517 | C15 | 99.67 | 2.00E-151 | JN558044.1 |
| seq525 | C15 | 99.33 | 8.00E-150 | JN558044.1 |
| seq526 | C15 | 99.67 | 6.00E-151 | JN558044.1 |
| seq527 | C15 | 99.66 | 2.00E-150 | JN558044.1 |
| seq532 | C15 | 99.33 | 8.00E-150 | JN558044.1 |
| seq543 | C15 | 99.33 | 8.00E-150 | JN558044.1 |
| seq548 | C15 | 99.33 | 3.00E-149 | JN558044.1 |
| seq565 | C15 | 99.67 | 6.00E-151 | JN558044.1 |
| seq569 | C15 | 99.33 | 8.00E-150 | JN558044.1 |
| seq570 | C3 | 99.67 | 2.00E-151 | MN876157.1 |
| seq572 | C15 | 99.67 | 2.00E-151 | JN558044.1 |
| seq579 | C15 | 99.67 | 6.00E-151 | JN558044.1 |
| seq591 | C15 | 99.33 | 8.00E-150 | JN558044.1 |
| seq592 | C15 | 99.67 | 6.00E-151 | JN558044.1 |
| seq603 | C15 | 99.33 | 8.00E-150 | JN558044.1 |
| seq605 | C15 | 99.33 | 8.00E-150 | JN558044.1 |
| seq613 | C15 | 99.67 | 2.00E-151 | JN558044.1 |
| seq614 | C3 | 99.67 | 2.00E-151 | MN876157.1 |
| seq633 | C15 | 99.33 | 8.00E-150 | JN558044.1 |
| seq638 | C15 | 99.33 | 8.00E-150 | JN558044.1 |
| seq655 | C15 | 99 | 4.00E-148 | JN558044.1 |
| seq676 | C15 | 99.33 | 8.00E-150 | JN558044.1 |
| seq679 | C15 | 99.33 | 8.00E-150 | JN558044.1 |
| seq692 | C15 | 99 | 1.00E-147 | JN558044.1 |
| seq697 | C15 | 99.33 | 8.00E-150 | JN558044.1 |
| seq710 | C15 | 99.33 | 3.00E-149 | JN558044.1 |
| seq722 | C15 | 99.33 | 8.00E-150 | JN558044.1 |
| seq726 | C15 | 98.33 | 8.00E-145 | JN558044.1 |
| seq731 | C15 | 99.67 | 6.00E-151 | JN558044.1 |
| seq749 | C15 | 99.33 | 3.00E-149 | JN558044.1 |
| seq752 | C15 | 99.67 | 2.00E-151 | JN558044.1 |
| seq756 | C15 | 99.33 | 3.00E-149 | JN558044.1 |
| seq760 | C15 | 99.33 | 8.00E-150 | JN558044.1 |
| seq765 | C15 | 99.33 | 3.00E-149 | JN558044.1 |
| seq770 | C15 | 99 | 4.00E-148 | JN558044.1 |
| seq776 | C15 | 99.67 | 2.00E-151 | JN558044.1 |
| seq784 | C1 | 99.67 | 2.00E-151 | AB778664.1 |
| seq805 | C15 | 98.66 | 6.00E-146 | JN558044.1 |
| seq807 | C15 | 99.67 | 6.00E-151 | JN558044.1 |
| seq814 | C15 | 99.33 | 8.00E-150 | JN558044.1 |
| seq824 | C15 | 99.33 | 8.00E-150 | JN558044.1 |
| seq848 | C15 | 99.33 | 8.00E-150 | JN558044.1 |
| seq850 | C15 | 98.33 | 8.00E-145 | JN558044.1 |
| seq862 | C15 | 99.33 | 4.00E-148 | JN558044.1 |
| seq873 | C15 | 99.67 | 2.00E-151 | JN558044.1 |
| seq877 | C15 | 99.67 | 2.00E-151 | JN558044.1 |
| seq898 | C15 | 99.33 | 8.00E-150 | JN558044.1 |
| seq913 | C15 | 99.33 | 8.00E-150 | JN558044.1 |
| seq947 | C15 | 99 | 1.00E-147 | JN558044.1 |
| seq953 | C91_ C15 | 99 | 4.00E-148 | C91: JN558050.1;  C15: JN558044.1 |
| seq962 | C91_C15 | 99 | 1.00E-147 | C91: JN558050.1;  C15: JN558044.1 |
| seq976 | C15 | 99.33 | 1.00E-148 | JN558044.1 |
| seq978 | C3 | 99 | 1.00E-147 | MN876157.1 |
| seq985 | C15 | 99.33 | 8.00E-150 | JN558044.1 |
| seq998 | C15 | 99.33 | 3.00E-149 | JN558044.1 |
| seq1014 | C3 | 99.67 | 2.00E-151 | MN876157.1 |
| seq1017 | C15 | 98.99 | 5.00E-147 | JN558044.1 |
| seq1025 | C15 | 99.33 | 8.00E-150 | JN558044.1 |
| seq1031 | C15 | 99.33 | 8.00E-150 | JN558044.1 |
| seq1032 | C15 | 99.33 | 8.00E-150 | JN558044.1 |
| seq1037 | C15 | 99.33 | 8.00E-150 | JN558044.1 |
| seq1046 | C3 | 99 | 4.00E-148 | MN876157.1 |
| seq1049 | C15 | 99.33 | 3.00E-149 | JN558044.1 |
| seq1060 | C15 | 99.33 | 3.00E-149 | JN558044.1 |
| seq1067 | C15 | 99.67 | 6.00E-151 | JN558044.1 |
| seq1070 | C15 | 99.67 | 2.00E-151 | JN558044.1 |
| seq1076 | C15 | 99.33 | 3.00E-149 | JN558044.1 |
| seq1085 | C15 | 99.33 | 4.00E-148 | JN558044.1 |
| seq1087 | C15 | 99.33 | 8.00E-150 | JN558044.1 |
| seq1097 | C15 | 99.33 | 8.00E-150 | JN558044.1 |
| seq1098 | C15 | 99.33 | 1.00E-148 | JN558044.1 |
| seq1105 | C15 | 99.33 | 3.00E-149 | JN558044.1 |
| seq1115 | C15 | 99 | 4.00E-148 | JN558044.1 |
| seq1130 | C15 | 99.66 | 2.00E-150 | JN558044.1 |
| seq1132 | C15 | 99.33 | 1.00E-148 | JN558044.1 |
| seq1139 | C15 | 99 | 1.00E-147 | JN558044.1 |
| seq1143 | C15 | 99.33 | 8.00E-150 | JN558044.1 |
| seq1144 | C15 | 98.67 | 2.00E-146 | JN558044.1 |
| seq1164 | C15 | 99.33 | 8.00E-150 | JN558044.1 |
| seq1169 | C15 | 98.99 | 5.00E-147 | JN558044.1 |
| seq1172 | C15 | 99 | 4.00E-148 | JN558044.1 |
| seq1173 | C15 | 99 | 1.00E-147 | JN558044.1 |
| seq1176 | C15 | 99.33 | 3.00E-149 | JN558044.1 |
| seq1178 | C15 | 99 | 4.00E-148 | JN558044.1 |
| seq1182 | C15 | 98.26 | 1.00E-137 | JN558044.1 |
| seq1184 | C15 | 99 | 1.00E-147 | JN558044.1 |

*Note:* There were 156 unclassified (NA) sequences of 1,185 ASVs curated in “DADA2”. Classification of NA sequences based on top hits from NCBI nt database using BLASTn program. For each query sequence, table includes sequence identification from “DADA2” output (ASV ID), classification (Genbank ID), the percent match between query and reference sequences (Percent Identity), the expect value (E value), and the accession number (Accession No.).

**Table S3. Data for fully analysed *P*. *lutea* and *P*. *lobata* samples (n = 37).**

| **Sample** | **No. of Reads** | **Site** | **Host** | **No. of Accum. Heat Stress Events** | **Taxa (*Cladocopium* ASVs)** | **No. of taxa** |
| --- | --- | --- | --- | --- | --- | --- |
| NING-MUI-POR-17 | 169971 | MUI | P. lobata | 3-6 | C15, C15.7, C15h, C60, C55, C15.9, C15.2.2, C15.8 | 8 |
| NING-MUI-POR-18 | 240618 | MUI | P. lobata | 3-6 | C15, C15h, C55, C15.9, C116, C56, C15.8, C15m, C15.6, C50 | 10 |
| NING-MUI-POR-20 | 208010 | MUI | P. lobata | 3-6 | C15, C15.7, C15h, C60, C55, C15.9, C15.2.2, C15.8 | 8 |
| NING-MUI-POR-10 | 225334 | MUI | P. lutea | 3-6 | C15, C15.7, C15h, C55, C15.9, C15g, C60, C15i, C15.8, C15.2.2 | 10 |
| NING-MUI-POR-14 | 190701 | MUI | P. lutea | 3-6 | C15, C15.9, C15.8, C15.7, C15h, C60, C15g, C15.2.2, C55, C56 | 10 |
| NING-MUI-POR-4 | 148773 | MUI | P. lutea | 3-6 | C15, C15.2.1, C116, C15h, C15.9, C15.7, C15j, C15i | 8 |
| NING-VLF-POR-7 | 165456 | VLF | P. lobata | 3-6 | C15, C116, C15.9, C55, C15.2.1, C15a, C15.6 | 7 |
| NING-VLF-POR-12 | 161108 | VLF | P. lutea | 3-6 | C15, C15L, C15h, C55, C56, C15.9, C15.8, C15m | 8 |
| NING-VLF-POR-13 | 169936 | VLF | P. lutea | 3-6 | C15, C15h, C56, C15a, C15m, C15.7, C15.6 | 7 |
| NING-VLF-POR-15 | 161917 | VLF | P. lutea | 3-6 | C15, C116, C15.9, C15.2.1, C15.6 | 5 |
| NING-VLF-POR-17 | 207084 | VLF | P. lutea | 3-6 | C15, C15h, C55, C56, C15.6, C15a, C15m | 7 |
| NING-VLF-POR-1 | 171663 | VLF | P. lutea | 3-6 | C15, C15h, C55, C15.6, C15m | 5 |
| NING-BUND-POR-10 | 110476 | BUN | P. lobata | 3-6 | C15, C15.1, C15L, C15.6 | 4 |
| NING-BUND-POR-2 | 207341 | BUN | P. lobata | 3-6 | C15, C116, C15a | 3 |
| NING-BUND-POR-4 | 152383 | BUN | P. lutea | 3-6 | C15, C116, C15h, C15.9, C55, C15.6 | 6 |
| NING-BUND-POR-6 | 173365 | BUN | P. lutea | 3-6 | C15, C116, C15h, C15.9, C15L, C15.7, C55 | 7 |
| NING-TAN-POR-4 | 199042 | TAN | P. lobata | 1-3 | C15, C55, C15f, C15.6, C91b | 5 |
| NING-TAN-POR-10 | 186107 | TAN | P. lutea | 1-3 | C15, C15h, C55, C15.9, C15.8, C15m, C15.6, C15.2.2 | 8 |
| NING-TAN-POR-11 | 169897 | TAN | P. lutea | 1-3 | C15, C116, C15.9, C55, C15.6, C15a, C15d, C15m, C15h | 9 |
| NING-TAN-POR-17 | 165767 | TAN | P. lutea | 1-3 | C15, C55, C15h, C15.6, C15c | 5 |
| NING-TAN-POR-1 | 170482 | TAN | P. lutea | 1-3 | C15, C15.9, C15.8, C15h, C60, C15.2.2, C15.7 | 7 |
| NING-TUR-POR-2 | 207040 | LK/TR | P. lobata | 1-3 | C15, C15.8, C15.7, C15h, C15.9, C60, C15.2.2, C15.6, C55 | 9 |
| NING-LAK-POR-1 | 160123 | LK/TR | P. lutea | 1-3 | C15, C15h, C56, C15.9, C15.8, C15m | 6 |
| NING-LAK-POR-2 | 159433 | LK/TR | P. lutea | 1-3 | C15, C56, C15.9, C15h, C15.8, C15m, C15.6, C15i | 8 |
| NING-LAK-POR-4 | 164964 | LK/TR | P. lutea | 1-3 | C15, C15.8, C15.7, C15h, C60, C15.9, C15.2.2, C15i | 8 |
| NING-LAK-POR-5 | 169603 | LK/TR | P. lutea | 1-3 | C15, C15.9, C15.8, C15.7, C15.2.2, C55, C15h | 7 |
| NING-TUR-POR-1 | 161517 | LK/TR | P. lutea | 1-3 | C15, C15h, C56, C15m, C15.6 | 5 |
| NING-OSP-POR-11 | 169498 | OSP | P. lobata | 1-3 | C15, C15f, C15.6, C15h, C91b | 5 |
| NING-OSP-POR-15 | 185777 | OSP | P. lobata | 1-3 | C15, C55, C15f | 3 |
| NING-OSP-POR-13 | 267804 | OSP | P. lutea | 1-3 | C15, C15m | 2 |
| NING-OSP-POR-2 | 241111 | OSP | P. lutea | 1-3 | C15, C15.9, C15.8, C15.7, C15h, C60, C55, C56, C15.2.2 | 9 |
| NING-OSP-POR-9 | 177571 | OSP | P. lutea | 1-3 | C15, C15.8, C15h, C60, C55, C15.9, C15g, C15.2.2, C15.7, C15.6 | 10 |
| NING-COR-POR-20 | 199036 | COR | P. lobata | 6-9 | C15, C15.9, C15.8, C15.7, C15h, C60, C15.2.2, C56 | 8 |
| NING-COR-POR-5 | 192191 | COR | P. lobata | 6-9 | C15, C116, C55, C15.6 | 4 |
| NING-COR-POR-9 | 198736 | COR | P. lobata | 6-9 | C15 | 1 |
| NING-COR-POR-13 | 210234 | COR | P. lutea | 6-9 | C15, C15.9, C15.7, C15h, C60, C15g, C116, C15.2.2, C91a, C15.8, C15.6, C56, C15d | 13 |
| NING-COR-POR-18 | 213593 | COR | P. lutea | 6-9 | C15, C15.9, C15.8, C15.7, C15h, C60, C55, C15.2.2, C15.6, C56 | 10 |

*Note:* Samples (rows) are listed in same order as shown in Figure 3.

**Table S4. Homogeneity of Dispersion Analysis (PERMDISP) Results (Formula: ~ Site).**

| **Factor** | **Df** | **SumOfSqs** | **MeanSq** | **F** | **p_value** |
| --- | --- | --- | --- | --- | --- |
| **Host** | **1** | **0.05606** | **0.05606** | **13.1129463** | **0.001** |
| Site | 6 | 0.02939281 | 0.0048988 | 0.32882609 | 0.917 |
| Temperature Range | 6 | 0.02939281 | 0.0048988 | 0.32882609 | 0.917 |
| No. of Accumulated Heat Stress Events | 2 | 0.00467456 | 0.00233728 | 0.24022494 | 0.787 |
| Warm Season Variability | 5 | 0.02357691 | 0.00471538 | 0.44002177 | 0.843 |
| Recovery Time | 3 | 0.0102062 | 0.00340207 | 0.35006667 | 0.794 |
| Max. Monthly Mean (MMM) | 5 | 0.02407804 | 0.00481561 | 0.339817 | 0.891 |

*Note:* Tests were run on variance stabilised data. Table includes a list of the main effects (Factor), degrees of freedom (Df), sum of squares (SumofSqs), mean squares (MeanSq), F statistic (F), and p-value (p_value), with all tests using n = 999 number of permutations. P-values of statistically significant Factors (p < 0.05) are **bold**.

**Table S5. PERMANOVA Results (Formula: ~ Site).**

| **Factor** | **Df** | **SumOfSqs** | **R_squared** | **pseudo_F** | **p_value** |
| --- | --- | --- | --- | --- | --- |
| **Host** | **1** | **0.2370074** | **0.05865809** | **2.18096434** | **0.007** |
| **Site** | **6** | **0.97310079** | **0.24083734** | **1.58620383** | **0.004** |
| Max. Monthly Mean (MMM) | 1 | 0.18404968 | 0.04555133 | 1.6703848 | 0.06 |
| **Temperature Range** | **1** | **0.26292002** | **0.06507133** | **2.43601086** | **0.002** |
| Warm Season Variability | 1 | 0.15563919 | 0.03851988 | 1.40220885 | 0.125 |
| **No. of Accum. Heat Stress Events** | **2** | **0.34830188** | **0.08620289** | **1.60369201** | **0.041** |
| Recovery Time | 1 | 0.09610434 | 0.02378532 | 0.85276957 | 0.585 |
| **Latitude** | **1** | **0.19681456** | **0.04871057** | **1.79216748** | **0.036** |
| Longitude | 1 | 0.14832578 | 0.03670985 | 1.33380878 | 0.16 |
| **Interactions** |  |  |  |  |  |
| Host:Site | 6 | 0.71541673 | 0.17706189 | 1.25903379 | 0.1 |
| Host:Temp. Range | 1 | 0.11464307 | 0.02837356 | 1.10399683 | 0.337 |
| Host:Warm Season Var. | 1 | 0.14964459 | 0.03703625 | 1.41364239 | 0.137 |
| Host:No. Heat Events | 2 | 0.27545959 | 0.0681748 | 1.3333638 | 0.124 |
| Host:Recovery | 1 | 0.13698975 | 0.03390424 | 1.26883156 | 0.188 |
| Host:Latitude | 1 | 0.12340707 | 0.0305426 | 1.16014585 | 0.276 |
| Host:Longitude | 1 | 0.12148442 | 0.03006676 | 1.13463697 | 0.294 |
| Host:MMM | 1 | 0.12325583 | 0.03050517 | 1.15541365 | 0.304 |
| **Temp. Range:Warm Season Var.** | **1** | **0.19212976** | **0.04755111** | **1.82212243** | **0.034** |
| Temp. Range:No. Heat Events | 1 | 0.12124853 | 0.03000838 | 1.16475036 | 0.28 |
| **Temp. Range:Recovery** | **1** | **0.20148262** | **0.04986589** | **1.91128416** | **0.029** |
| Temp. Range:MMM | 1 | 0.06735922 | 0.01667105 | 0.63721983 | 0.869 |
| Temp. Range:Latitude | 1 | 0.10319717 | 0.02554076 | 0.98640403 | 0.427 |
| **Temp. Range:Longitude** | **1** | **0.19866609** | **0.04916882** | **1.89218973** | **0.025** |
| **No. Heat Events:Warm Season Var.** | **1** | **0.20322104** | **0.05029614** | **1.89945715** | **0.031** |
| No. Heat Events:Recovery | 1 | 0.13787773 | 0.03412402 | 1.32063913 | 0.189 |
| **No. Heat Events:Latitude** | **2** | **0.54692812** | **0.13536184** | **2.78019015** | **0.001** |
| **No. Heat Events:Longitude** | **2** | **0.48641709** | **0.12038568** | **2.44507092** | **0.001** |
| No. Heat Events:MMM | 1 | 0.1616431 | 0.04000582 | 1.49203971 | 0.112 |
| **Warm Season Var.:Recovery** | **1** | **0.23658334** | **0.05855314** | **2.26802316** | **0.009** |
| Warm Season Var.:Latitude | 1 | 0.09333266 | 0.02309934 | 0.85540293 | 0.597 |
| Warm Season Var.:Longitude | 1 | 0.08886204 | 0.02199289 | 0.80107706 | 0.669 |
| Warm Season Var.:MMM | 1 | 0.10425222 | 0.02580188 | 0.95759081 | 0.498 |
| Recovery:Latitude | 1 | 0.15362999 | 0.03802262 | 1.40526832 | 0.113 |
| **Recovery:Longitude** | **1** | **0.26034641** | **0.06443437** | **2.44299855** | **0.006** |
| Recovery:MMM | 1 | 0.12934381 | 0.03201192 | 1.17166713 | 0.248 |

*Note:* The table includes a list of the main effects and interactions (Factor), degrees of freedom (Df), sum of squares (SumOfSqs), coefficient of determination (R_squared), pseudo-F statistic (pseudo_F), and p-value (p-value). P-values of statistically significant Factors (p < 0.05) are **bold**.

**Table S6. Mean Alpha Diversity across Categorical Factors.**

| **Factor** | **Group** | **Mean_Shannon** | **SE_Shannon** | **Mean_Simpson** | **SE_Simpson** |
| --- | --- | --- | --- | --- | --- |
| Host | P. lobata | 2.12095429 | 0.1362063 | 0.77455845 | 0.0246777 |
|  | P. lutea | 2.27205915 | 0.0860431 | 0.77183151 | 0.0175363 |
| Site | MUI | 2.44347529 | 0.1699008 | 0.80246642 | 0.0342344 |
|  | VLF | 2.22883735 | 0.2396186 | 0.76286229 | 0.0525035 |
|  | BUN | 2.13094244 | 0.3697659 | 0.7931773 | 0.0759384 |
|  | TAN | 2.19907283 | 0.2130064 | 0.76398128 | 0.0421521 |
|  | LK/TR | 2.18037821 | 0.1333006 | 0.74718161 | 0.0252996 |
|  | OSP | 2.24126341 | 0.1235109 | 0.77925319 | 0.0151331 |
|  | COR | 2.05204588 | 0.1886551 | 0.76585454 | 0.0175078 |
| No. of Accumulated | 1-3 | 2.2052469 | 0.0855373 | 0.76245387 | 0.0160910 |
| Heat Stress Events | 3-6 | 2.28485285 | 0.1368358 | 0.78529259 | 0.0282913 |
|  | 6-9 | 2.05204588 | 0.1886551 | 0.76585454 | 0.0175078 |

**Table S7. Table of Shapiro-Wilk Test Results.**

| **Metric** | **W** | **p_value** |
| --- | --- | --- |
| Shannon | 0.98561929 | 0.90461614 |
| Simpson | 0.98093513 | 0.76354136 |

*Note:* Checking assumption of normality for alpha diversity metrics.

**Table S8. Levene’s Test Results for Alpha Diversity.**

| **Factor** | **Metric** | **Df** | **F** | **p_value** |
| --- | --- | --- | --- | --- |
| Host | Shannon | 1 | 0.26469436 | 0.61014695 |
|  | Simpson | 1 | 0.10049665 | 0.75311776 |
| Site | Shannon | 6 | 0.52378712 | 0.78565398 |
|  | Simpson | 6 | 0.96236726 | 0.46702041 |
| No. of Accumulated Heat Stress Events | Shannon | 2 | 0.72921722 | 0.48967635 |
|  | **Simpson** | **2** | **4.54950997** | **0.0177499** |

*Note:* Checking assumption of Homogeneity of Variances for alpha diversity metrics. Statistically significant values (*p* < 0.05) are **bold**.

**Table S9. Independent *t*-test Results for Alpha Diversity.**

| **Factor** | **Metric** | **t_value** | **df** | **conf_low** | **conf_high** | **p_value** |
| --- | --- | --- | --- | --- | --- | --- |
| Host | Shannon | -0.9379144 | 21.6865647 | -0.4855012 | 0.18329145 | 0.35861744 |
|  | Simpson | 0.09007542 | 23.987877 | -0.059757 | 0.06521092 | 0.92897518 |

**Table S10. Welch’s ANOVA Results for Alpha Diversity.**

| **Factor** | **Metric** | **F_value** | **num_df** | **denom_df** | **p_value** |
| --- | --- | --- | --- | --- | --- |
| Site | Shannon | 0.35634043 | 6 | 12.6112189 | 0.89329472 |
|  | Simpson | 0.29632812 | 6 | 12.5833659 | 0.92758287 |
| No. of Accumulated Heat Stress Events | Shannon | 0.47102752 | 2 | 11.2478369 | 0.63613864 |
|  | Simpson | 0.24124334 | 2 | 16.5877356 | 0.78835924 |

**Table S11. Summary of Distribution of ASVs and *Cladocopium* taxa across categorical factors.**

| **Factor** |  | **# of ASVs** | ***Cladocopium* taxa** | **# of taxa** |
| --- | --- | --- | --- | --- |
| **Site** | MUI | 285 | C15, C15h, C116, C56, C15.2.2, C55, C60, C15.9, C15.2.1, C15.8, C15m, C15i, C15.7, C15.6, C50, C15g, C15j | 17 |
|  | VLF | 284 | C15, C15.9, C55, C15h, C15.2.1, C15a, C116, C15.6, C56, C15m, C15.8, C15L, C15.7 | 13 |
|  | BUN | 110 | C15, C15.9, C116, C15.7, C15L, C15h, C15a, C15.6, C55, C15.1 | 10 |
|  | TAN | 321 | C15, C15h, C116, C15.9, C55, C15d, C15.6, C15.8, C15m, C15.7, C15.2.2, C91b, C15c, C60, C15a, C15f | 16 |
|  | LK/TR | 290 | C15, C15h, C15.6, C15.9, C15.8, C60, C15m, C15.7, C56, C15.2.2, C55, C15i | 12 |
|  | OSP | 234 | C15, C15h, C56, C60, C15.9, C15.8, C15.6, C15m, C15.7, C15.2.2, C55, C15g, C15f, C91b | 14 |
|  | COR | 143 | C15h, C15, C56, C15d, C15.9, C15.8, C15.6, C91a, C116, C60, C15.7, C15.2.2, C55, C15g | 14 |
| **Host Species** | P. lobata | 424 | C15, C15h, C116, C56, C15.9, C55, C60, C15a, C15.6, C15.8, C15.2.1, C15m, C15.7, C15.2.2, C91b, C15L, C15.1, C50, C15f | 19 |
|  | P. lutea | 783 | C15, C15h, C116, C56, C15.9, C15.2.2, C15.6, C55, C15d, C60, C15.2.1, C15.7, C15.8, C91a, C15L, C15m, C15i, C15a, C15c, C15g, C15j | 21 |
| **No. of Accumulated Heat Stress Events** | 1-3 | 623 | C56, C15, C15.9, C55, C15h, C116, C15d, C15.6, C15a, C15.8, C15c, C15g, C60, C91b, C15i, C15f, C15.7, C15.2.2, C15m | 19 |
|  | 3-6 | 542 | C15.6, C116, C15, C15.9, C15L, C56, C50, C15.2.2, C15h, C15a, C55, C15.8, C15.7, C15.2.1, C15.1, C15j, C15g, C60, C15i, C15m | 20 |
|  | 6-9 | 143 | C15, C15d, C116, C56, C91a, C15h, C15.8, C55, C15.9, C60, C15.6, C15g, C15.7, C15.2.2 | 14 |

**Table S12. Prevalence and relative abundance of each *Cladocopium* taxon across full dataset (n = 37 samples)**

| ***Cladocopium* taxon** | **Total number of ASVs mapping to taxon** | **Prevalence (number of samples)** | **Abundance per sample (mean)** | **Abundance per sample (SE)** |
| --- | --- | --- | --- | --- |
| C15 | 708 | 37 | 0.89121928 | 0.0163268 |
| C15h | 35 | 28 | 0.00711777 | 0.00076168 |
| C15.9 | 29 | 24 | 0.01442997 | 0.00183848 |
| C55 | 16 | 22 | 0.00326422 | 0.00086089 |
| C15.6 | 45 | 20 | 0.00377889 | 0.00145067 |
| C15.8 | 13 | 18 | 0.00930289 | 0.00098057 |
| C15.7 | 7 | 16 | 0.00862791 | 0.00094272 |
| C15.2.2 | 9 | 14 | 0.07261286 | 0.00819011 |
| C56 | 26 | 12 | 0.01280746 | 0.00574061 |
| C60 | 7 | 12 | 0.00325453 | 0.00039725 |
| C15m | 5 | 11 | 0.00662946 | 0.00149805 |
| C116 | 49 | 10 | 0.12615454 | 0.02962209 |
| C15a | 8 | 5 | 0.00603009 | 0.00366164 |
| C15g | 2 | 4 | 0.00271617 | 0.00080676 |
| C15i | 7 | 4 | 0.003716 | 0.00160511 |
| C15.2.1 | 6 | 3 | 0.10518867 | 0.05501074 |
| C15L | 8 | 3 | 0.01034702 | 0.00576469 |
| C15f | 2 | 3 | 0.00493456 | 0.00080215 |
| C15d | 7 | 2 | 0.00344845 | 0.00062321 |
| C91b | 4 | 2 | 0.00493355 | 0.00148218 |
| C15.1 | 2 | 1 | 0.03879576 | NA |
| C15c | 2 | 1 | 0.00215966 | NA |
| C15j | 1 | 1 | 0.00252062 | NA |
| C50 | 1 | 1 | 0.00012468 | NA |
| C91a | 2 | 1 | 0.0049326 | NA |

**Table S13. Frequency and proportion of ASVs, mapping to *Cladocopium* taxa, across each heat stress category**

| **Category** | **Taxon** | **Frequency** | **Fraction** |
| --- | --- | --- | --- |
| **low (1-3 events) - all** | C116 | 15 | 0.024077 |
|  | C15 | 459 | 0.736758 |
|  | C15.2.2 | 6 | 0.009631 |
|  | C15.6 | 34 | 0.054575 |
|  | C15.7 | 3 | 0.004815 |
|  | C15.8 | 9 | 0.014446 |
|  | C15.9 | 14 | 0.022472 |
|  | C15a | 3 | 0.004815 |
|  | C15c | 2 | 0.00321 |
|  | C15d | 3 | 0.004815 |
|  | C15f | 2 | 0.00321 |
|  | C15g | 2 | 0.00321 |
|  | C15h | 24 | 0.038523 |
|  | C15i | 2 | 0.00321 |
|  | C15m | 5 | 0.00802568 |
|  | C55 | 11 | 0.0176565 |
|  | C56 | 18 | 0.02889246 |
|  | C60 | 7 | 0.01123596 |
|  | C91b | 4 | 0.00642055 |
| **moderate (3-6 events) - all** | C116 | 35 | 0.064576 |
|  | C15 | 377 | 0.695572 |
|  | C15.1 | 2 | 0.00369 |
|  | C15.2.1 | 6 | 0.01107 |
|  | C15.2.2 | 6 | 0.01107 |
|  | C15.6 | 10 | 0.01845 |
|  | C15.7 | 7 | 0.012915 |
|  | C15.8 | 7 | 0.012915 |
|  | C15.9 | 20 | 0.0369 |
|  | C15a | 8 | 0.01476 |
|  | C15g | 2 | 0.00369 |
|  | C15h | 21 | 0.038745 |
|  | C15i | 5 | 0.009225 |
|  | C15j | 1 | 0.001845 |
|  | C15L | 8 | 0.01476 |
|  | C15m | 2 | 0.00369 |
|  | C50 | 1 | 0.001845 |
|  | C55 | 8 | 0.01476 |
|  | C56 | 12 | 0.02214 |
|  | C60 | 4 | 0.00738 |
| **High (6-9 events) - all** | C116 | 10 | 0.06993007 |
|  | C15 | 90 | 0.62937063 |
|  | C15.2.2 | 3 | 0.02097902 |
|  | C15.6 | 2 | 0.01398601 |
|  | C15.7 | 3 | 0.02097902 |
|  | C15.8 | 6 | 0.04195804 |
|  | C15.9 | 5 | 0.03496504 |
|  | C15d | 4 | 0.02797203 |
|  | C15g | 1 | 0.00699301 |
|  | C15h | 8 | 0.05594406 |
|  | C55 | 3 | 0.02097902 |
|  | C56 | 3 | 0.02097902 |
|  | C60 | 3 | 0.02097902 |
|  | C91a | 2 | 0.01398601 |

**Supplementary Figures**

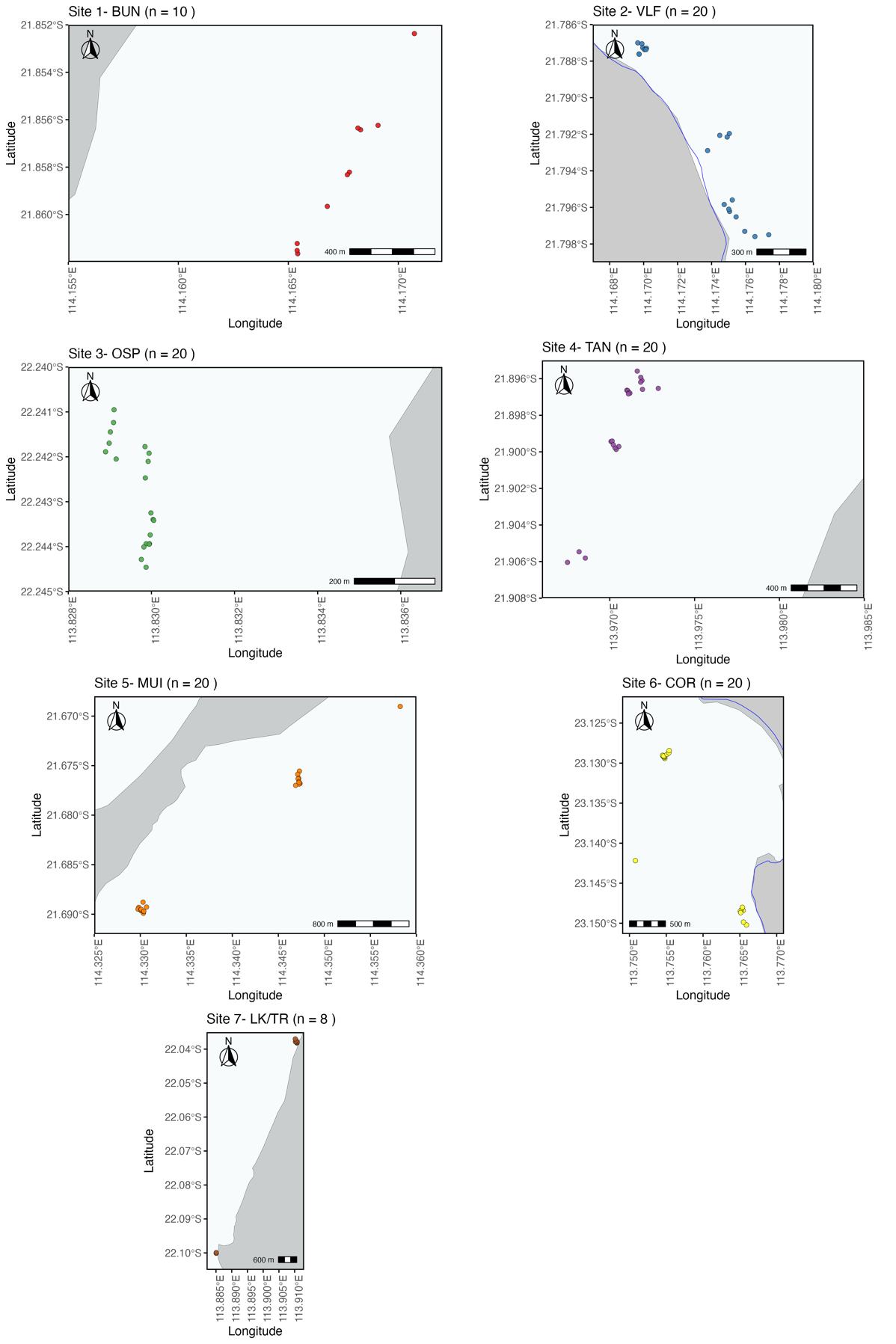

**Figure S1. Site Maps.** Each panel indicates site number, site name (abbreviated), and number of samples collected at that site (n = X). Points represent GPS coordinates of all sampled *Porites* colonies per site and are colored according to site (see Fig. 1). Additional colours represent land (grey), water (light blue), and UNESCO Ningaloo World Heritage Coast boundary line (dark blue). Google Earth (<https://earth.google.com>) was used to generate KML files incorporating GPS coordinates of sampled *Porites* colonies and land boundaries within the study area.

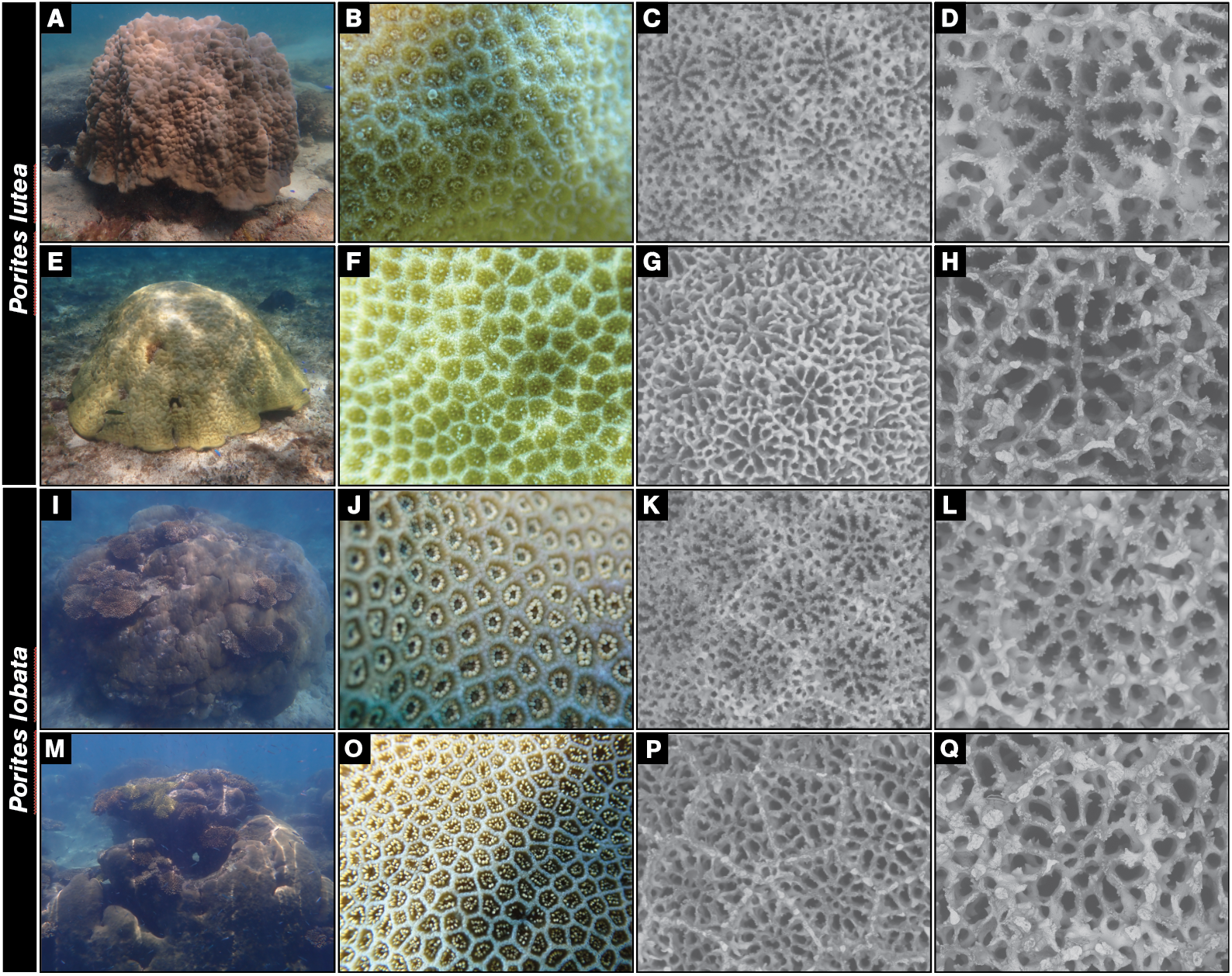

**Figure S2. A-D *P*. *lutea* classic (S2-P.15); E-H *P*. *lutea* lightly calcified (S2-P.12); I-L *P*. *lobata* classic (S4_P.05); M-Q *P*. *lobata* (S4-P.14) lightly calcified. Each row corresponds to one individual colony.**

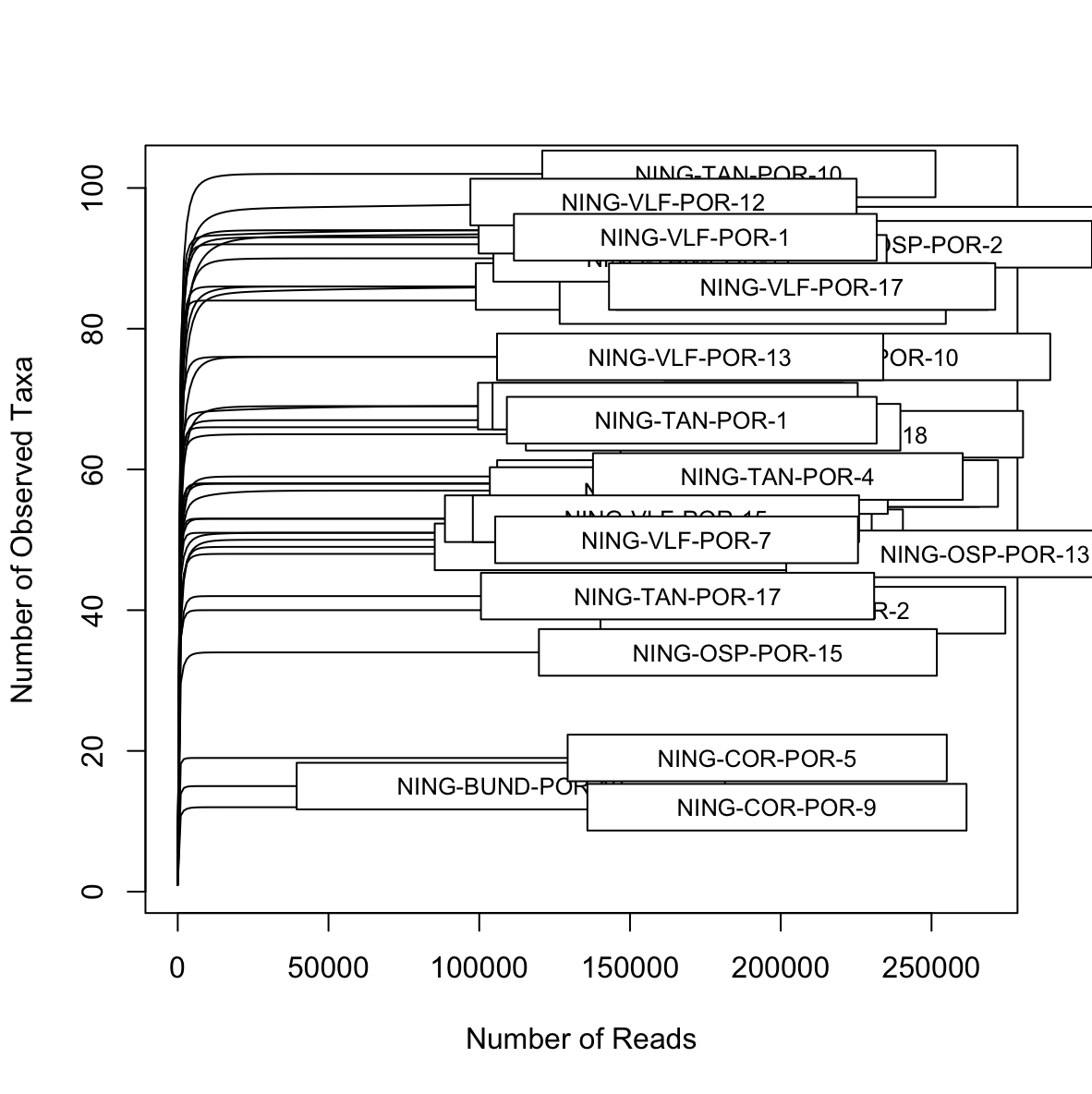

**Figure S3. Rarefaction Curves of Untransformed *P*. *lobata* and *P*. *lutea* Samples (n = 37).**

**(a)	(b)**

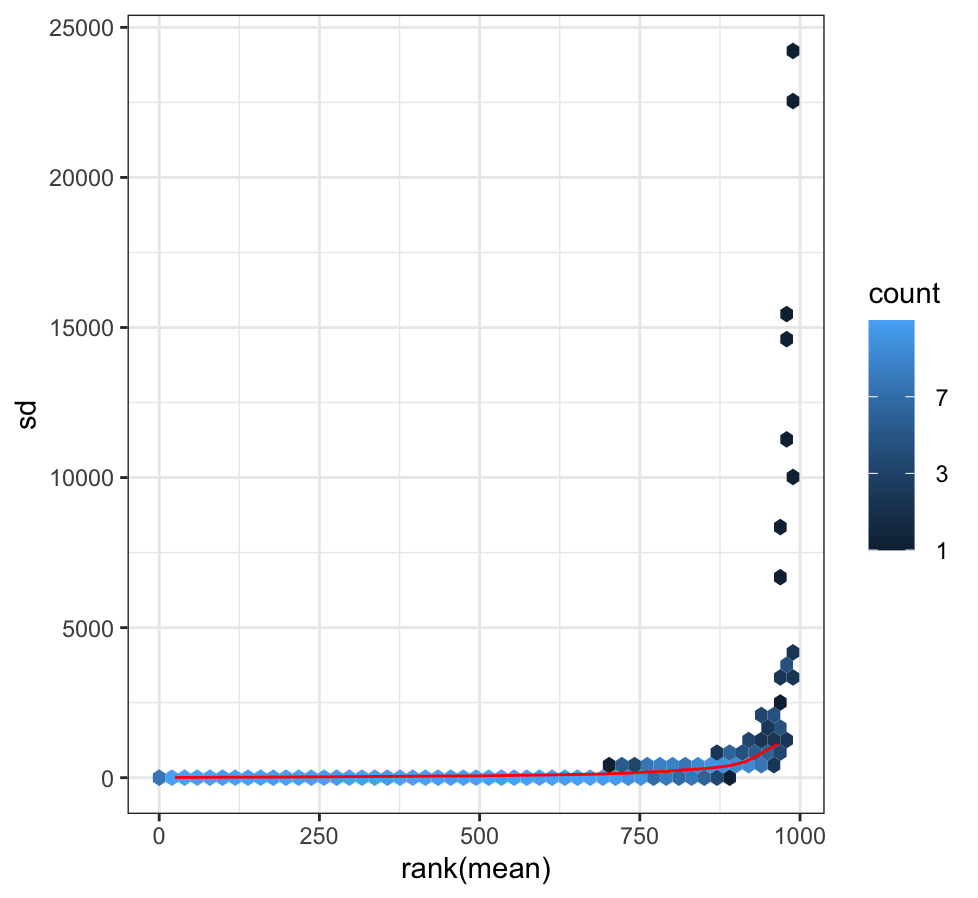

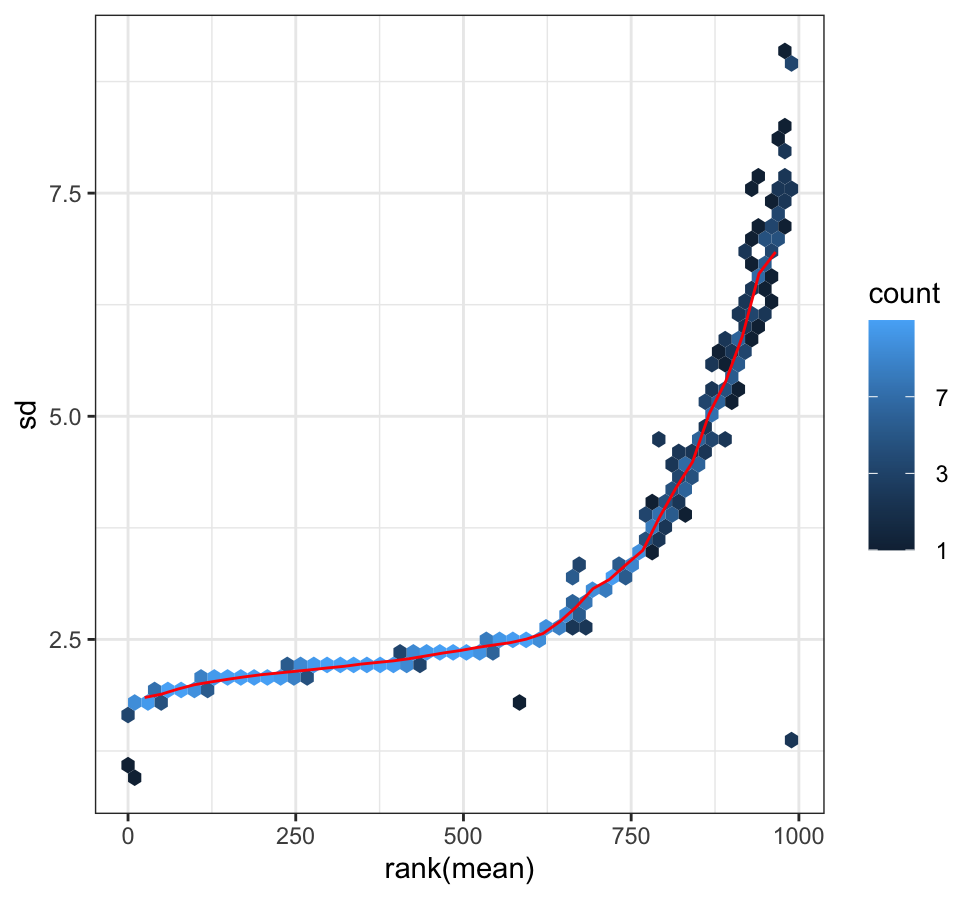

**(c)	(d)**

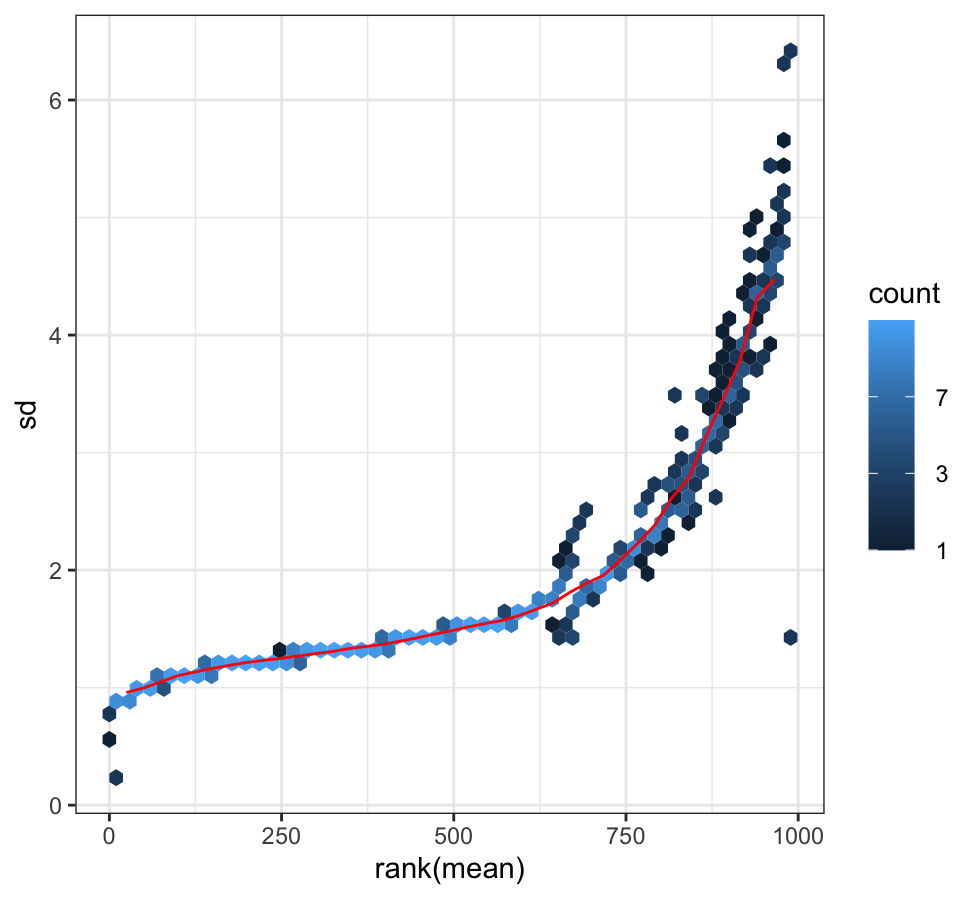

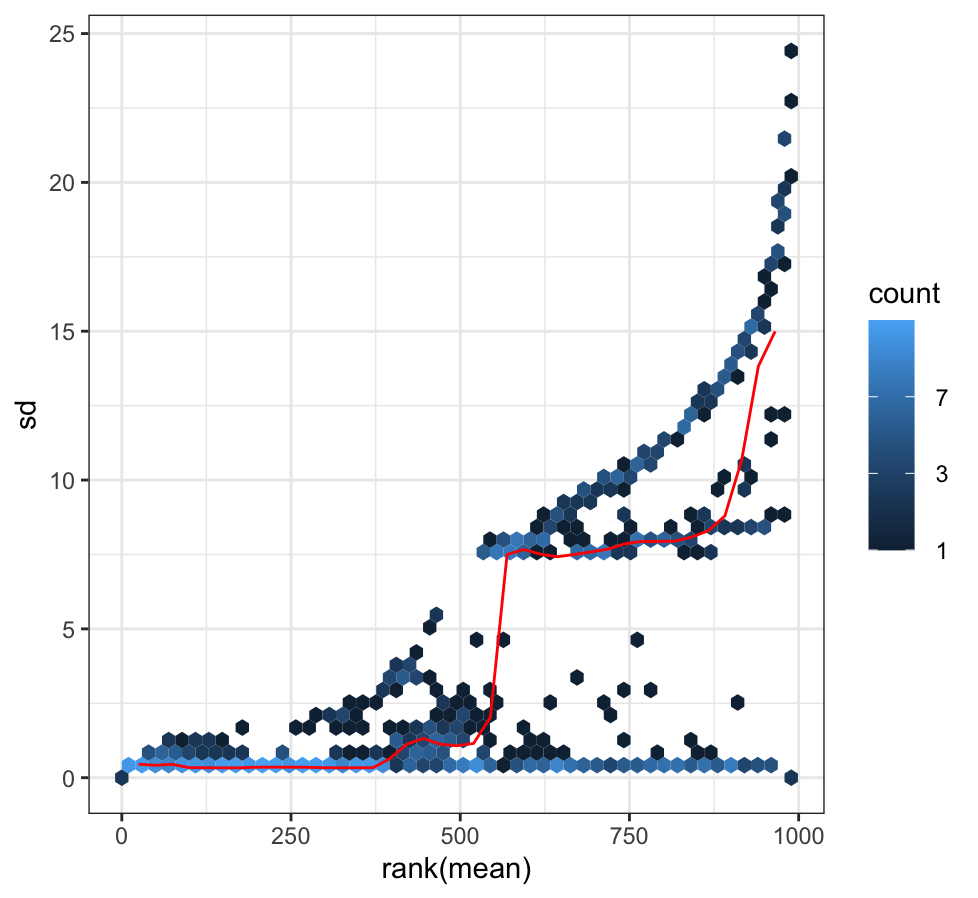

**Figure S4. Standard Deviation Plots of Data Transformations.** Plots of the standard deviation of the data, across samples, against the mean for: (a) untransformed data, (b) variance stabilised transformed data, (c) shifted logarithm transformed (log2(n + 1)), and (d) regularized log transformed data.

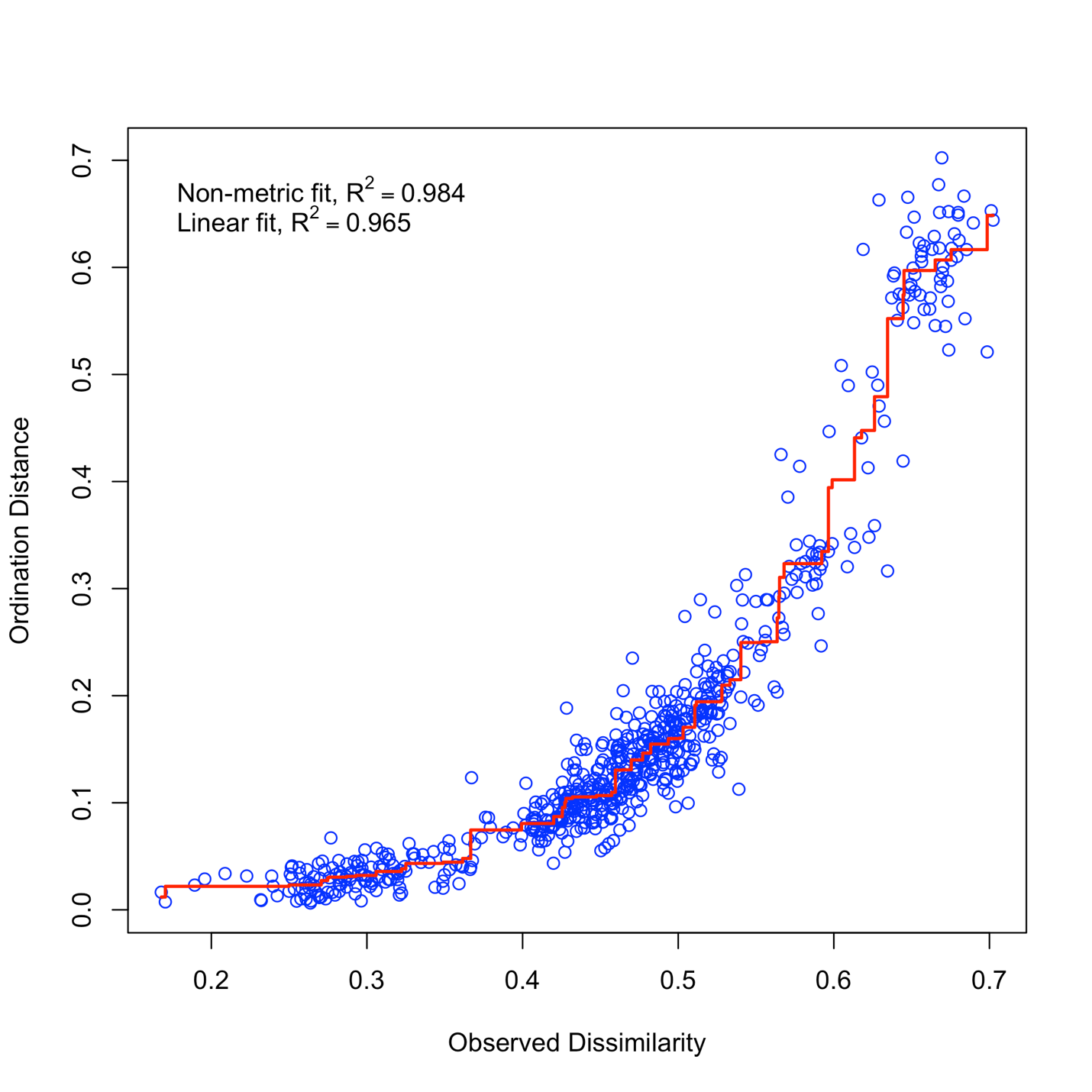

**Figure S5. Stress Plot for Weighted UniFrac Dissimilarity.** The plot evaluates the goodness of fit of the NMDS ordination (stress = 0.1258899), showing the relationship between the observed dissimilarity (x-axis) and the ordination distance (y-axis). The non-metric fit (R² = 0.984) and the linear fit (R² = 0.965) indicate a high level of accuracy in representing the data in reduced dimensions.

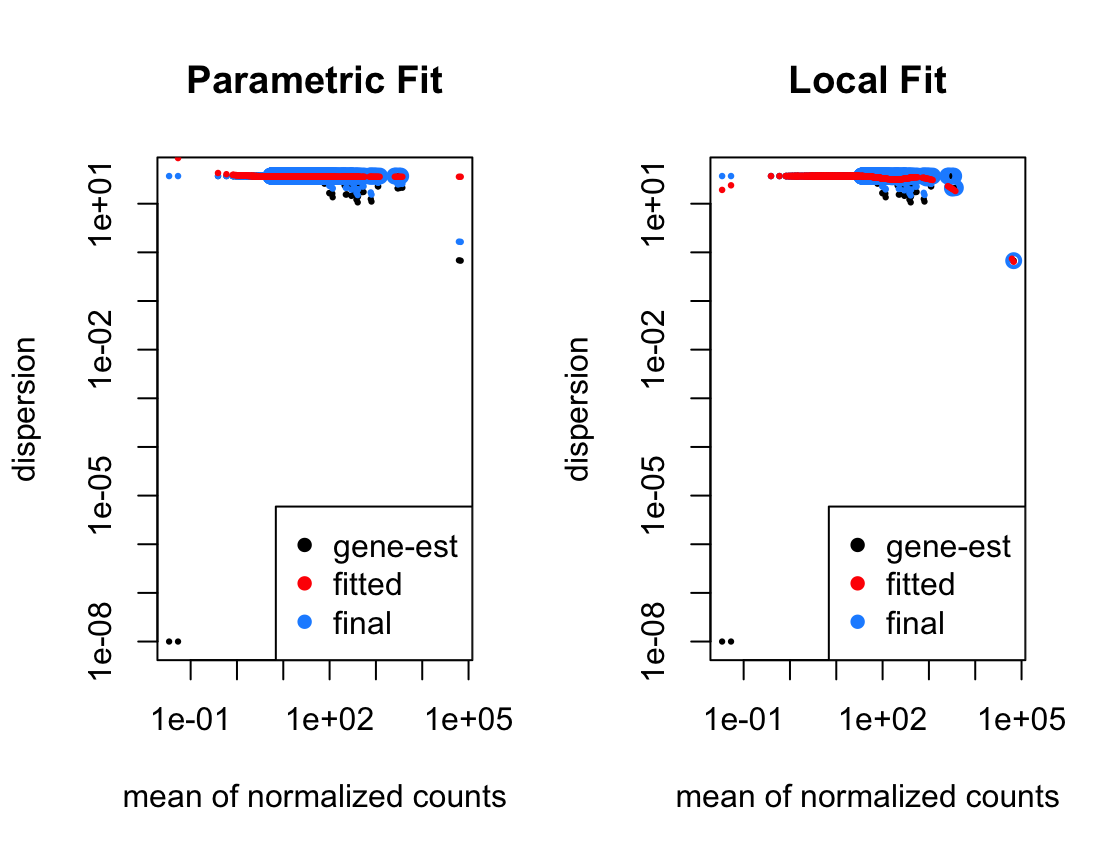

**Figure S6.** **Dispersion Estimate Plots of Various Fit Types.** Dispersion estimates were tested using (a) “parametric” and (b) “local” fitting of dispersions (fit types) to the mean of normalised counts.

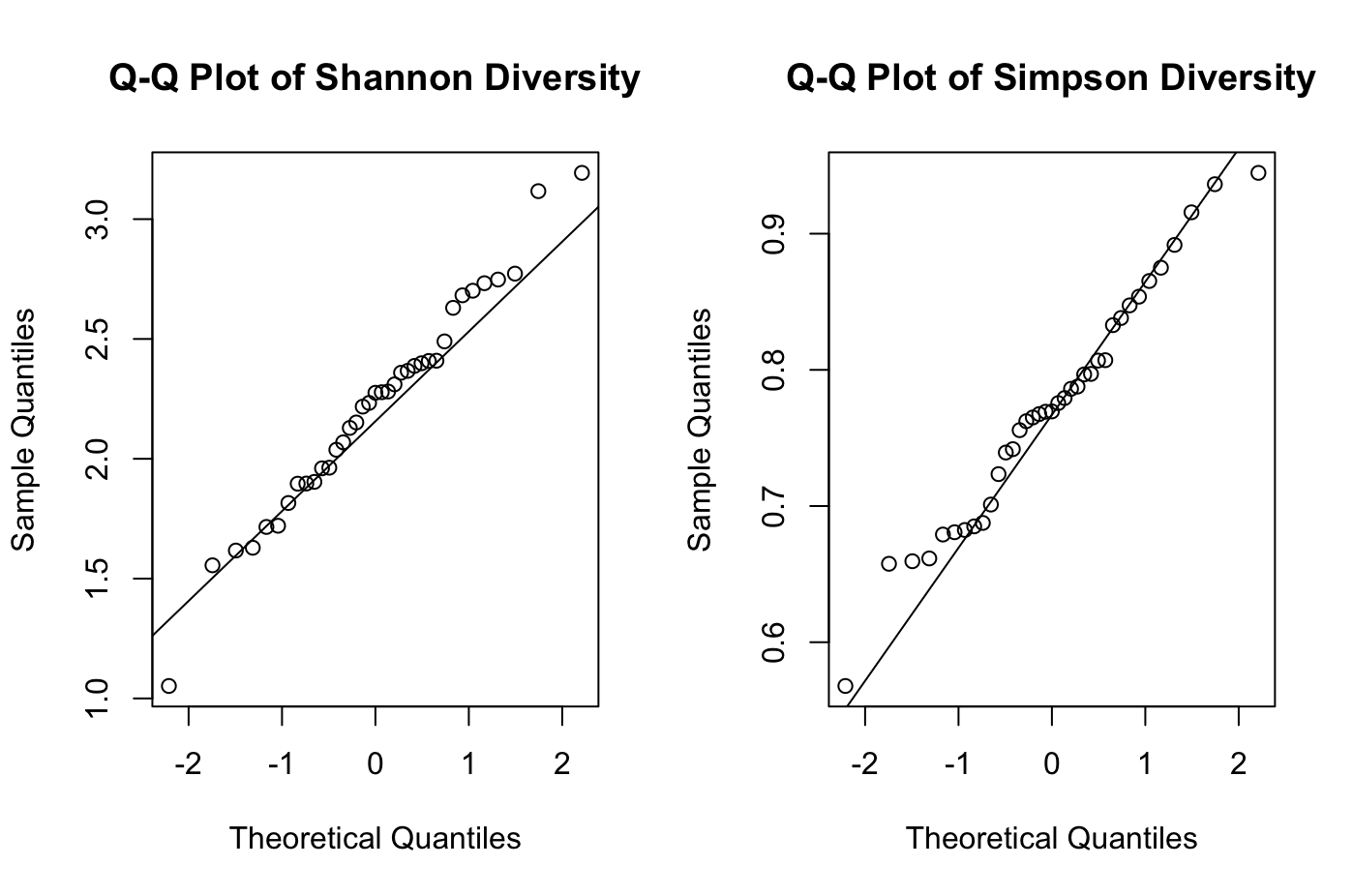

**Fig S7. Q-Q Plots of Alpha Diversity – visual test for normal distribution.**

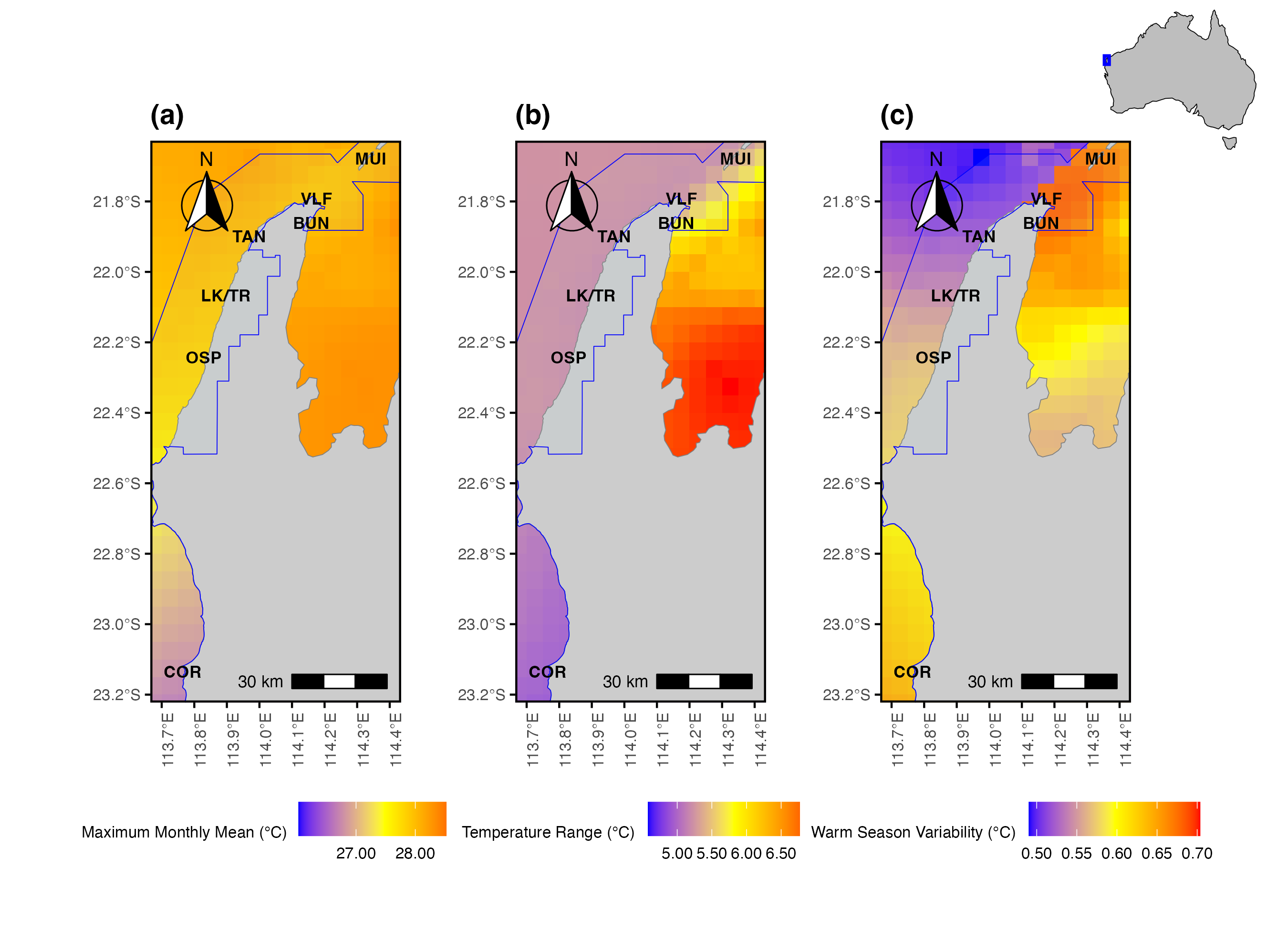

**Fig S8. Temperature-related environmental variables across sites extracted from NOAA Coral Reef Watch NetCDF4 data.** Maps display significant temperature-related variables derived from 5 km-resolution satellite time series products (NOAA Coral Reef Watch, 2024), with missing or non-analyzed areas predicted using Kriging interpolation. Blue outlines indicate the boundary of UNESCO World Heritage Ningaloo Coast, with site names labeled. **(a)** Maximum Monthly Mean (MMM) sea surface temperatures (°C) **(b)** annual temperature range (°C) **(c)** warm season variability (°C). The inset map (top right) shows the study region along the Ningaloo Reef, Western Australia (blue square).

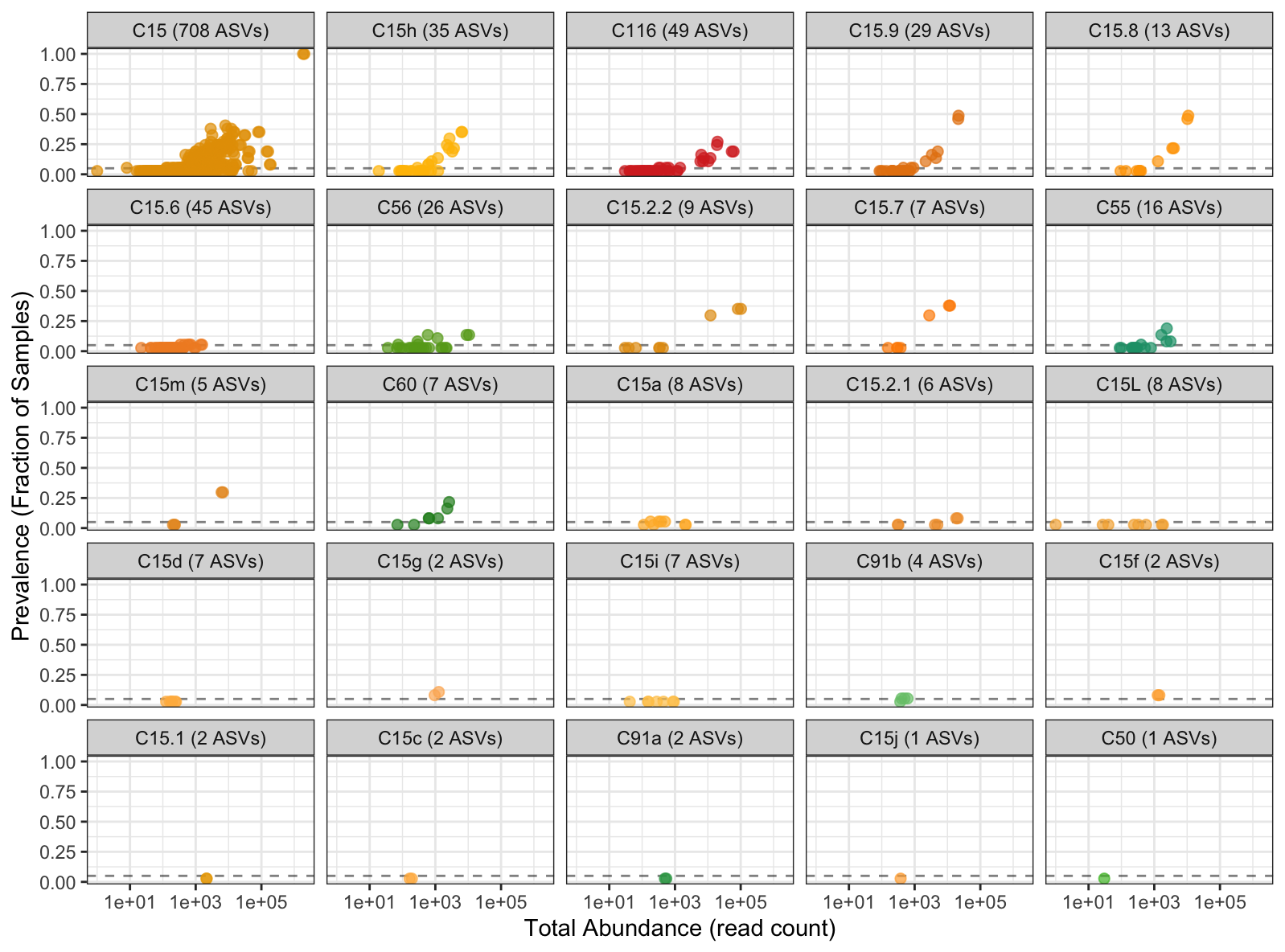

**Figure S9. Cumulative Symbiodiniaceae Prevalence across Samples.** ASVs are represented by dots, coloured according to the *Cladocopium* taxon to which they map to. The plot is facetted by *Cladocopium* taxa observed across the cumulative 37 *P*. *lutea* and *P*. *lobata* samples. The position of dots represent the relationship between the prevalence of each ASV across samples (y-axis) and the abundance of ASVs within the read count (x-axis). Facet titles include the number of ASVs (out of 1,001 total) that map to each *Cladocopium* taxa. Facets are ordered by most prevalent to least (left to right, top to bottom).

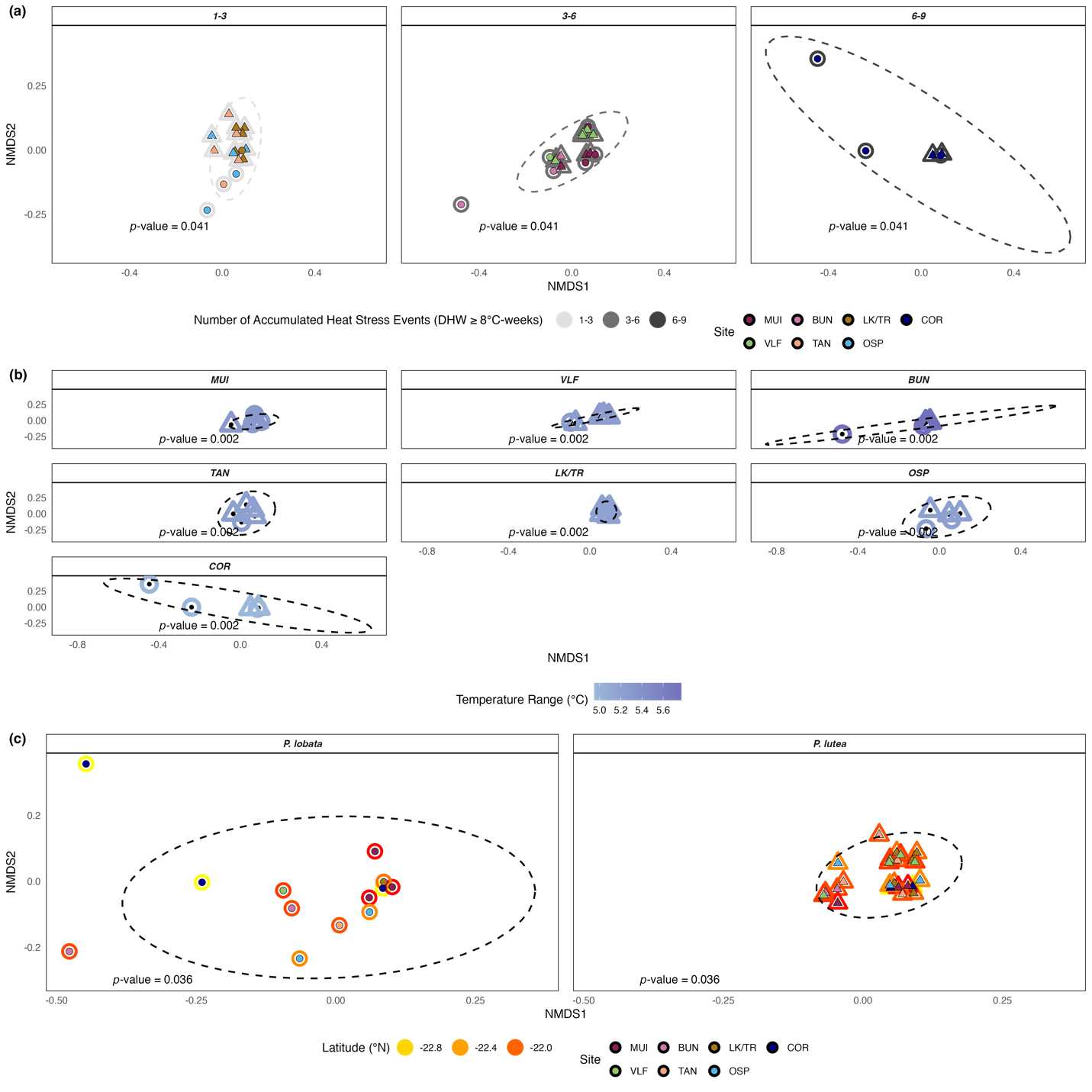

**Figure S10.** Non-metric multidimensional scaling (nMDS) ordinations of Symbiodiniaceae community composition based on weighted UniFrac distances (stress = 0.126). Points represent unique Symbiodiniaceae communities associated with individual *P*. *lobata* (circle) and *P*. *lutea* (triangle) samples. Dashed ellipses show 95% confidence intervals for group clustering. The significant PERMANOVA p-value associated with each represented factor (color) is noted at the bottom left of each NMDS plot. **(a)** Ordinations grouped and coloured by the number of heat stress events (DHW ≥ 8°C-WEEKS; 1–3 events, 3–6 events, 6–9 events), with site designated by inner color. **(b)** Ordinations grouped by sites, colors represent mean temperature range, **(c)** Ordinations for *P*. *lobata* (left) and *P*. *lutea* (right), colored by latitude and highlighting significant differences in dispersion between the two hosts (PERMDISP: *F* = 13.113, *p* = 0.001).

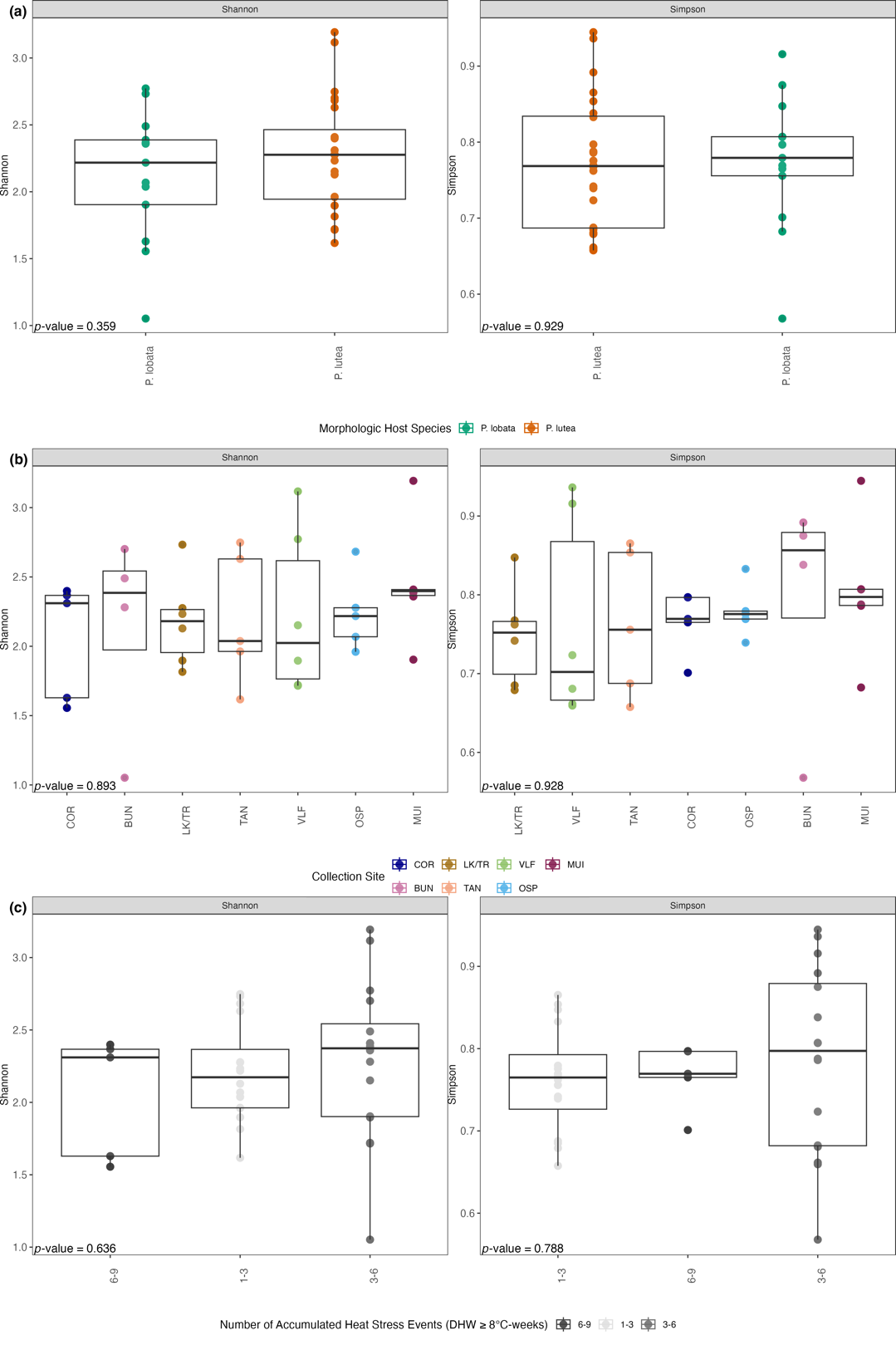

**Figure S11.** **Boxplots of Alpha Diversity Metrics for Different Factors.** Points represent the diversity value (y-axis) for each sample within each group (x-axis), the black box shows the interquartile range (IQR) where the middle 50% of the data points lie; median diversity is represented by the black bar. Groups are ordered from least to greatest mean diversity (left to right). Factors are as follows: **(a)** morphologic host species, **(b)** sites, **(c)** number of accumulated heat stress events (DHW ≥ 8°C-weeks).

**Supplementary Data**

**Data S1. DNA Extraction Protocol.**

Kit: QIAGEN DNeasy Blood & Tissue Kit

Protocol: Purification of Total DNA from Animal Tissues (Spin-Column Protocol) with **minor modifications** as follows:

1. Approximately **20 mg** (+/- 3mg) of tissue was scraped off coral skeleton with a scalpel and immediately placed into sterile 1.7 ml microcentrifuge tubes containing 180 µl Buffer ATL.
2. 20 µl Proteinase K was then added to each sample. Samples were pulse-vortexed for 10-15 sec to initiate chemical lysis.
3. Samples underwent **incubation at 56°C for 16 hours** using a Hybridisation Oven set to level 3.
4. Following heat lysis in oven, samples were pulse-vortexed for 15 seconds. 200 µl Buffer AL was added to each sample and then pulse-vortexed again for 10 seconds. Immediately after this, 200 µl ethanol absolute was added to each sample and then pulse-vortexed again for 15 seconds.
5. Clear mixture was pipetted out of the original microcentrifuge tubes into labelled DNeasy Mini spin column placed in 2 ml collection tubes. Caution was used to avoid transfer of coral skeleton and excess mucus.
6. Samples were centrifuged for 1 minute at 6000 x g (rcf); **additional centrifugation (same speed, same duration) was performed for complete filtration as necessary**. Flow-through and collection tubes were discarded.
7. DNeasy Mini spin column were placed in new collection tubes. 500 µl Buffer AW1 was added to each column and centrifuged for 1 minute at 6000 x g (rcf); **additional centrifugation (same speed, same duration) was performed for complete filtration as necessary**. Flow-through and collection tubes were discarded.
8. DNeasy Mini spin column were placed in new collection tubes and 500 µl Buffer AW2 and centrifuged for 3 minutes at 20,000 x g (rcf) to dry DNeasy membrane. Caution was used when discarding flow-through and collection tube to ensure no residual ethanol was carried over into the elution step. If residual ethanol was still present, centrifugation for 1 minute at 20,000 x g (rcf) was required before moving on to the next step.
9. DNeasy Mini spin column was placed in a sterile, labelled 1.7 ml microcentrifuge. Per protocol recommendation - elution was performed **in two rounds**. For each round, **75 µl Buffer AE** was pipetted directly onto DNeasy membrane and incubated at room temperature for 1 minute before centrifugation for 1 minute at 6000 x g (rcf). The same 1.7 ml microcentrifuge tube was used for both rounds of elution. The total elute volume was 150 ml.
10. Elutes were placed on ice until Nanodrop quantification.
11. Following quantification, elutes were stored in -20°C freezer until PCR amplification.

**Data S2. Standard PCR protocol using MyTaq DNA Polymerase (Bioline Inc.)**

PCR reaction set-up was prepared on ice in a PCR Cabinet (Esco Lifesciences Group). Each 20 µl PCR included 4 µl of 5x My Taq reaction buffer, 0.8 µl each 0.4µM Forward (5’-GTGAATTGCAGAACTCCGTG-3’) and Reverse (5’-CCTCCGCTTACTTATATGCTT-3’) primers, 0.2 µl of 2.5U MyTaq DNA polymerase, 13.2 µl of MilliQ water, and 1 µl of DNA template. PCR amplification cycles were performed on an Eppendorf Mastercycler^®^ nexus GSX1 as follows: 95°C for 3 min; followed by 35 cycles at 95°C for 30 s, 59°C for 30 s, 72°C for 30 s, and 72°C for 7 min. PCR products were visualized on 1.5% agarose gels. For PCR product that did not produce a visible line using initial DNA template, dilutions of DNA template were performed, and PCR was redone until a consistent, visible line was confirmed by gel electrophoresis.
